# Erlotinib induces 3D genome rearrangements in lung cancer cells activating tumor suppressor genes through FOXA2-bound Epromoters

**DOI:** 10.1101/2025.05.13.653883

**Authors:** Guruprasadh Swaminathan, Julio Cordero, Stefan Günther, Johannes Graumann, Thomas Braun, Gergana Dobreva, Guillermo Barreto

## Abstract

Non-small cell lung cancer (NSCLC) is the most frequent lung cancer (LC). While erlotinib is an epidermal growth factor (EGFR) tyrosine kinase inhibitor (TKI) used in the treatment of NSCLC, it remains unclear how this FDA-approved drug affects the genome. We performed integrative multi-omics studies in human pulmonary carcinoma cells to elucidate the epigenetic mechanisms induced by erlotinib. We identified 746 genes (including 34 tumor suppressor genes, TSG) that were upregulated after treatment with erlotinib or gefitinib (another EGFR-TKI). Interestingly, 45% of the upregulated genes (including 24 TSG) were in broad domains of the euchromatin histone mark H3K4me3, and 63% (including 26 TSG) exhibited reduced levels of the heterochromatin histone mark H3K27me3 after erlotinib treament. Further, H3K27ac-specific chromosome conformation capture-based methods revealed that erlotinib significantly increased number and length of chromatin loops between promoters of upregulated genes and active enhancers. We also detected augmented chromatin accessibility after erlotinib treatment at the promoters of upregulated genes, which correlated with binding of the transcription activator FOXA2. Remarkably, we identified gene clusters that seem to be upregulated by promoters with enhancer activity (Epromoters) enriched with FOXA2. The clinical relevance of our findings was confirmed by data from The Cancer Genome Atlas, showing significantly improved survival outcomes in LC patients with high levels of *FOXA2* and/or the 34 TSG found upregulated by erlotinib. Our results establish 3D genome rearrangements as molecular mechanism mediating EGFR-TKI effects in NSCLC cells, supporting the design of more specific therapies for NSCLC targeting different chromatin features.

## INTRODUCTION

Genomic alterations in tumor suppressor genes (TSG, such as *TP53* and *RB1*) and oncogenes (such as *EGFR, KRAS, ALK, RET* and *ROS1*) are known to drive malignant transformation in cancer [1–5]. However, while some cancers develop because of genomic alterations, others do not, suggesting alternative events that trigger cancer. Specific chromatin rearrangements provide a plausible explanation for those cancers that arise without somatic mutations, since changes in chromatin are key events that must occur before cancer-related genes are expressed [6–8]. Eukaryotic transcription occurs within the physiological template of chromatin, which is hierarchically organized at different levels including chromosomal territories, compartments, self-interacting topologically associating domains, and chromatin loops, altogether resulting in a highly dynamic three-dimensional (3D) genome organization that plays an important role in transcriptional regulation [9]. Chromatin can be structurally condensed blocking the access of the transcription machinery to transcriptionally inactive regions (heterochromatin), or structurally relaxed mediating the access of the transcription machinery to transcriptionally active regions (euchromatin). Post-translational histone modifications are one of the mechanisms by which chromatin structure and transcription are regulated. Tri-methylated lysine 4 of histone 3 (H3K4me3) is a well-characterized euchromatin histone mark related to genes with high transcriptional activity [10,11]. Additionally, studies have demonstrated that genes with broad H3K4me3 domains exhibit higher transcriptional activity compared to those with narrow domains [11,12]. Moreover, H3K4me3 broad domains are associated with increased transcription elongation and enhancer activity, which together lead to high expression of TSG [11]. On the other hand, tri-methylated lysine 27 of histone 3 (H3K27me3) is a well-characterized heterochromatin histone mark that is deposited by the poly-comb repressive complex 2 (PRC2) and is associated with transcriptional repression for cell type-specific genes [13–16]. Further, H3K27me3 is characteristic for distal regulatory elements in the genome that are capable of silencing gene expression (so called silencers), whereas it also accumulates over intergenic regions and non-transcribed gene bodies forming large blocks of H3K27me3-marked loci [17–20].

Structurally less-condensed euchromatin not only facilitates access of the transcriptional machinery to promoters, but also binding of transcription factors (TFs) and co-activators to typical enhancers (TYE), which are relatively short (∼100–1000 bp) DNA sequences that function as distal regulatory elements controlling transcription of their cognate promoters [21–23]. In addition, super-enhancers (SE) are clusters of enhancers enriched with specific histone marks (such as histone 3 monomethylated at lysine 4 or acetylated at lysine 27, H3K4me1 and H3K27ac respectively), cofactors (such as components of the multimeric protein complex cohesin and mediator of RNA polymerase II transcription subunit 1, MED1) and cell-type-specific TFs [24–26]. Interestingly, Epromoters have been reported as promoters with structural and functional characteristics of enhancers regulating not only the expression of their cognate gene but also of other distal genes [27–30]. Chromatin loops can bring into close physical proximity two distant sequences of DNA. Thus, chromatin looping is broadly accepted as a means for enhancer-promoter interactions [31].

Pioneer TFs belonging to the FOXA protein family are embryonic master regulators playing a crucial role in the development of organs that arise from the endoderm, such as the lung, pancreas, liver, colon and prostate [32–36]. Accumulating evidence supports that FOXA pioneer TFs induce embryonic expression signatures in non-cancerous somatic cells causing malignant transformation during cancer initiation [7,37]. Forkhead Box A2 (FOXA2) is an embryonic master regulator from the FOXA family of pioneer TFs that has been related to a range of cancer subtypes [36,38–42]. In the context of lung cancer (LC), FOXA2 functions as a TSG, playing a critical role in preserving epithelial integrity and halting cancer progression [43,44]. FOXA2 can bind its target DNA sequence at promoters or enhancers even within compacted chromatin and mediate changes in chromatin structure that allow binding of other TFs to induce cell-type-specific gene signatures maintained through mitotic cell division [45]. The pioneered opening of regulatory elements at facultative heterochromatin and the subsequent binding of TFs mediating signal-dependent, cell-type-specific transcription is referred to as assisted loading [45]. Interestingly, pioneer factors exhibit cell-type-specific binding patterns and can be excluded from specific chromatin structures such as constitutive heterochromatin [46,47].

LC is histologically classified into small cell lung cancer (SCLC) and non-small cell lung cancer (NSCLC) [48]. Whereas SCLC is a highly aggressive cancer that spreads very fast and represents 15% of all LC cases [2,49], NSCLC progresses at a comparatively slower rate and accounts for 85% of all LC cases [50]. NSCLC is further subcategorized into lung adenocarcinoma (LUAD; 45%), squamous cell carcinoma (LUSC; 25%), and large cell carcinoma (15%) [51]. Despite advancements in treatments, LC remains the leading cause of cancer-related dead worldwide [52], and early detection remains essential for improving survival rates [53–56]. Activation of epidermal growth factor receptor (EGFR) plays a key role in LC progression [57,58]. Mutations in the EGFR are identified in 32% of NSCLC cases, predominantly in LUAD. The majority of these EGFR mutants have either multi-nucleotide in-frame deletions in exon 19 (ΔEx19), or a point mutation in exon 21 of *EGFR*, in which leucine 858 is substituted by arginine (L858R), each of these somatic mutations resulting in activation of the tyrosine kinase domain of EGFR. The small molecules erlotinib and gefitinib are EGFR tyrosine kinase inhibitors (TKIs) used to manage and treat specific NSCLC and pancreatic cancer [59,60]. LUADs responsive to these EGFR-TKI possess the *EGFR* mutations described above and often increased *EGFR* copy number [61,62]. Erlotinib and gefitinib inhibit EGFR tyrosine kinase activity by reversibly competing with ATP for binding in the kinase domain. However, the effects on the genome of LC cells caused by EGFR-TKI have been sparsely studied. Here, we performed integrative multi-omics studiesto determine the effect of blocking EGFR by erlotinib on the 3D genome of NSCLC cells to elucidate the epigenetic mechanisms leading to expression of TSG, thereby providing further insights into the therapeutic effects of EGFR-TKIs, as well as the molecular basis for designing more specific therapies for NSCLC targeting different chromatin features.

## RESULTS

### Erlotinib increases H3K4me3 broad domains associated with gene activation

We analyzed the transcriptome of a human alveolar basal epithelial adenocarcinoma cell line (A549) after erlotinib treatment by total RNA sequencing (RNA-seq, Fig. 1a-b, Supplementary Fig. 1a-b, Source Data file). From the transcripts that were significantly affected by erlotinib treatment (n = 1,002), a minority (n = 256; 25.5%) showed reduced expression after erlotinib treatment, whereas 74.5% (n = 746; further referred to as upregulated transcripts or genes) showed increased expression with a median of −0.6 log2 RPKM+1 and an interquartile range (IQR) of 3.9 log2 RPKM+1 (P = 3.49E-283), as compared to −1.37 log2 RPKM+1 (IQR = 3.86 log2 RPKM+1) in control (Ctrl) non-treated cells (Fig. 1b, top). These 746 upregulated genes after erlotinib treatment included 34 genes that were previously reported as TSG (Source Data file). Further, we also detected significantly increased expression of the 746 upregulated genes in four different publicly available RNA-seq datasets (Supplementary Fig. 1c, Source Data file), including erlotinib treated A549 cells [63], as well as PC-9 cells treated with erlotinib, gefitinib or osimertinib [64–66], confirming our results in two human pulmonary adenocarcinoma cell lines treated with different EGFR-TKIs. Interestingly, single-cell RNA-seq (scRNA-seq) in a lung adenocarcinoma patient-derived xenograft tumor model, in which tumor-bearing mice were non-treated or erlotinib treated [67] showed that the cell cluster with highest *EGFR* levels was the only cell cluster with significantly increased expression of the 34 TSG included in the 746 upregulated genes (Supplementary Fig. 1d). These scRNA-seq results in a lung adenocarcinoma patient-derived xenograft tumor model confirmed the RNA-seq results in two human pulmonary adenocarcinoma cell lines (Fig. 1a-b; Supplementary Fig. 1c), as well as demonstrated a direct correlation between *EGFR* levels and cell sensitivity to erlotinib. To determine the clinical relevance of the *in vitro* RNA-seq and *in vivo* scRNA-seq results, we analyzed RNA-seq data from control donors and NSCLC patients deposited at The Cancer Genome Atlas (TCGA; Fig. 1c). We found that expression levels of the 746 upregulated genes were significantly lower in non-treated NSCLC patients (median = 2.06 log2 TPM+1; IQR = 4.16 log2 TPM+1; P = 5.53E-5) as compared to Ctrl donors (median = 2.2 log2 TPM+1; IQR = 5.12 log2 TPM+1). In addition, expression levels of the 746 upregulated genes significantly increased in NSCLC patients that were treated with erlotinib (median = 3.23 log2 TPM+1; IQR = 4.49 log2 TPM+1; P = 3.35E-110) as compared to non-treated NSCLC patients. Similar results were obtained by analyzing the RNA-seq data from control donors and NSCLC patients deposited at TCGA focusing on the expression of 34 TSG included in the 746 upregulated genes (Fig. 1c, bottom; Source Data file). These findings in Ctrl donors and NSCLC patients non-treated or treated with erlotinib confirmed the results in human pulmonary adenocarcinoma cell lines (Fig. 1a-b; Supplementary Fig. 1c) and in the lung adenocarcinoma patient-derived xenograft tumor model (Supplementary Fig. 1d). Gene Set Enrichment Analysis (GSEA) of the differentially expressed genes (DEGs) after erlotinib treatment (Fig. 1d) revealed significant enrichment of signaling pathways related to cancer, including EGFR-resistance, PI3K-AKT-, mTOR-, MAPK-, TGFB- and NFκB-signaling across the upregulated transcripts, whereas glycolysis, WNT-, TNF- and IL-17-signaling were significantly enriched in the down-regulated transcripts. To further investigate the effect of erlotinib treatment in lung adenocarcinoma cells, we performed a sequencing experiment following chromatin immunoprecipitation (ChIP-seq) for genome-wide profiling of H3K4me3 (Fig. 1e-h, Supplementary Fig. 2a, Source Data file). Correlating with the effect of erlotinib treatment in the transcriptome of A549 cells, we detected genome-wide increase of H3K4me3 levels after erlotinib treatment (Fig. 1e, top; median = 0.47 RPMM; IQR = 3.31E-3 RPMM; P = 1.17E-26), as compared to Ctrl non-treated cells (median = 0.44 RPMM; IQR = 6.51E-3 RPMM). Since the increasing effect of erlotinib in H3K4me3 levels was not observed at transcription start sites (TSS) -/+2 kb (Fig. 1e, bottom), we performed genome-wide peak distribution analysis of the H3K4me3 ChIP-seq and found that the number of H3K4me3 peaks increased after erlotinib treatment at promoters and gene bodies as compared to Ctrl non-treated cells (Fig. 1f). To gain further insight into this finding, we determined broad and narrow domains of H3K4me3 (Fig. 1g) and found that H3K4me3 broad domains spanning promoters and gene bodies showed a significant increase in erlotinib-treated cells from 79.5% to 85.1% (P < 1E-4) for promoters and from 25.3% to 27.3% (P = 0.03) for gene bodies, compared to control non-treated cells, whereas non-significant change was observed in intergenic regions. Further analysis showed that 338 genes coding for transcripts that were increased after erlotinib treatment as determined by RNA-seq in two different human pulmonary adenocarcinoma cell lines (Fig. 1a-b, Supplementary Fig. 1c) were located in H3K4me3 broad domains and included 26 known TSG (Fig. 1h; Supplementary 2b). Focusing on the H3K4me3 broad domains spanning these 338 upregulated genes (Fig. 1i, top; Source Data file) we detected significantly increased H3K4me3 levels after erlotinib treatment at promoters (median = 2.37 RPMM; IQR = 1.28 RPMM; P = 9.9E-12) and gene bodies (median = 1.93 RPMM; IQR = 0.89 RPMM; P = 4.1E-7), as compared to the same promoters (median = 2.12 RPMM; IQR = 1.09 RPMM) and gene bodies (median = 1.67 RPMM; IQR = 0.55 RPMM) in Ctrl non-treated cells. Focusing on H3K4me3 broad domains spanning promoters and bodies of genes that were downregulated after erlotinib treatment (Fig. 1i, bottom), erlotinib treatment did not significantly change H3K4me3 levels. The levels of H3K4me3 significantly increased after erlotinib treatment at promoters and gene bodies of these 338 genes that were upregulated after erlotinib treatment, supporting that H3K4me3 broad domains are associated with increased transcription elongation.

**Figure 1:**
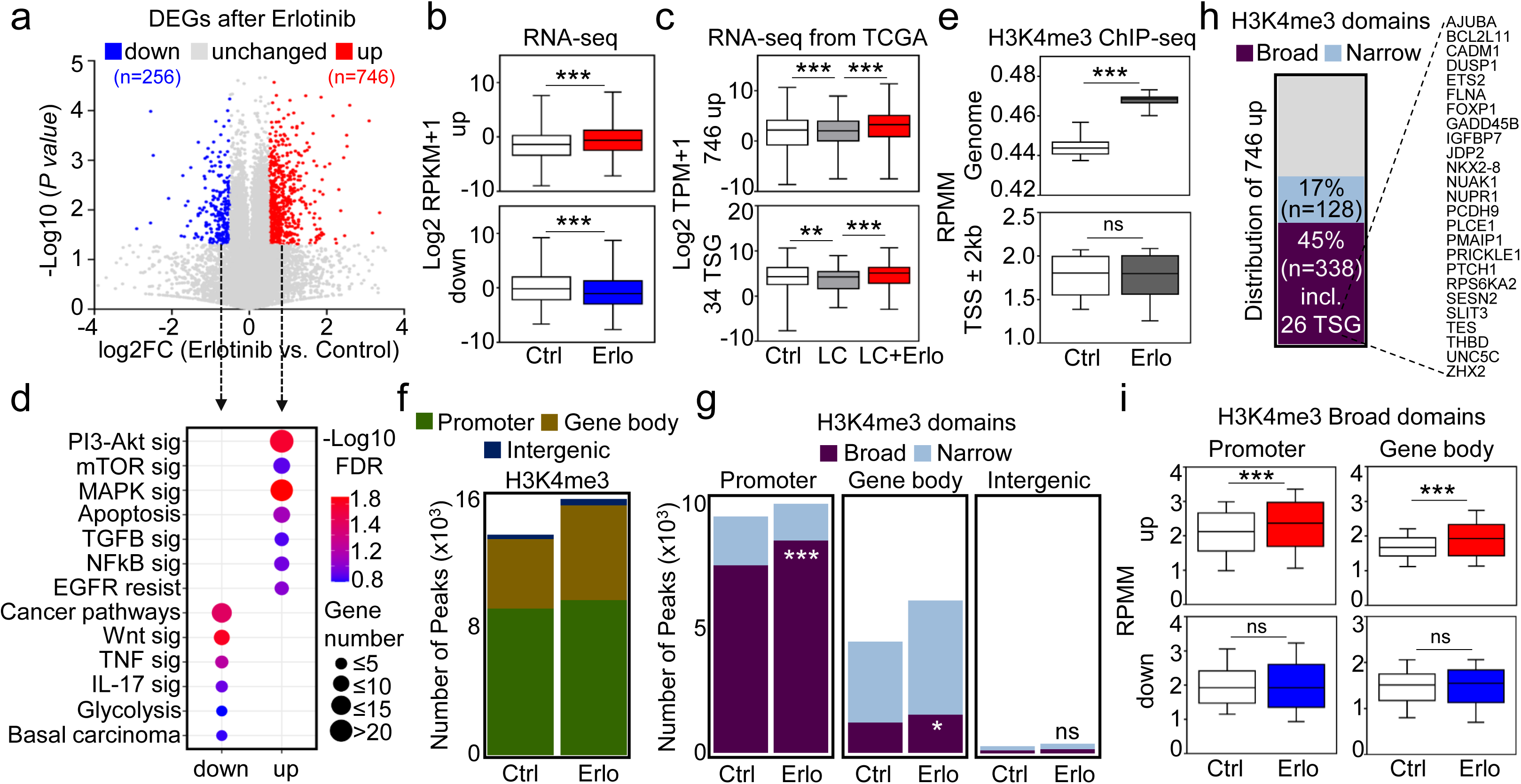
Erlotinib increases H3K4me3 broad domains associated with gene activation. (**a**) RNA-sequencing in human lung adenocarcinoma cells. A549 cells were treated with control (DMSO) or 5 µM erlotinib for 48 h. Volcano plot represents the significance (-log10 *P*-values after paired two-tailed t-test) vs. expression log2 fold change (log2FC ratios) between the average of control (Ctrl) and erlotinib (Erlo) treated cells. Differentially expressed genes (DEGs) of upregulated (up; n=746) and downregulated (down; n=256) genes were identified by a cutoff of *P* < 0.05. (**b**) Box plots of RNA-seq-based expression analysis of DEGs after erlotinib. (**c**) Box plots of RNA-seq-based expression analysis from TCGA of upregulated (up; n=746) and tumor suppressor genes (TSG; n=34) in normal lung (Ctrl; n=59), patients with non-small cell lung cancer (LC; n=1,050), and patients with lung adenocarcinoma treated with erlotinib (LC+Erlo; n=22). (**d**) Gene set enrichment analysis (GSEA) of upregulated and downregulated genes as identified in **1a**. sig, signaling; EGFR resist, EGFR resistance. FDR, false discovery rate. (**e**) Box plots showing the levels of H3K4me3 in Ctrl or Erlo treated A549 cells genome-wide or at transcription start site (TSS -/+ 2 kb). Bar plots showing the distribution of H3K4me3 peaks (**f**) and bar plots displaying the broadness of H3K4me3 (**g**) in different genomic regions at Promoters (Peaks -/+ 2 kb from TSS), Gene body (exon and intron regions outside the -/+ 2 kb TSS), and Intergenic (peaks not located in previous regions) in Ctrl or Erlo treated A549 cells. (**h**) Bar plot with the distribution of upregulated genes in the H3K4me3 domains (45% broad; n=338, including 26 TSG, 17% narrow; n=128). (**i**) Box plots showing the levels of H3K4me3 in the significantly upregulated or downregulated genes that were separated into the indicated H3K4me3 domains (broad or narrow), and genomic regions (promoter or gene body). In all box plots, values were normalized using RPKM, reads per kilobase of transcript per million mapped reads or TPM, transcript per million; represented as log2 RPKM + 1 or log2 TPM + 1. All box plots display the median (middle line), 25th and 75th percentiles (box), and 5th and 95th percentiles (whiskers). Statistical significance is represented by asterisks: \*\*\**P* ≤ 0.001; \*\**P* ≤ 0.01; \**P* ≤ 0.05; ns, non-significant. *P*-values were calculated after two-tailed t-test (box plots) or two-tailed Fisher exact test (bar plots). See also Supplementary Fig. 1 and Supplementary Fig. 2. Source data are provided as a Source Data file.

### Gene activation induced by erlotinib correlates with reduced H3K27me3 broad domains

To further investigate the effect of erlotinib on the chromatin landscape, we performed a sequencing experiment following Cleavage Under Targets and Tagmentation (CUT&Tag) for high-resolution genome-wide profiling of the heterochromatin histone mark H3K27me3 in Ctrl non-treated or erlotinib treated A549 cells (Fig. 2a-d, Supplementary Fig. 3a-c; Source Data file). We detected significant genome-wide reduction of H3K27me3 levels after erlotinib treatment (Fig.2a, top; median = 0.98 RPMM; IQR = 0.08 RPMM; P = 9.32E-13) as compared to Ctrl non-treated cells (median = 1 RPMM; IQR = 0.1 RPMM). In contrast to the results obtained for H3K4me3 at the TSS -/+2kb (Fig. 1e, bottom), we detected significant reduction of H3K27me3 levels at TSS -/+2kb after erlotinib treatment (Fig. 2a, bottom). However, in reflection of our observations for H3K4me3, genome-wide peak distribution analysis of the H3K27me3 CUT&Tag revealed that the majority of the H3K27me3 peaks were also in broad domains (Supplementary Fig. 3b). In addition, H3K27me3 broad domains spanning promoters and gene bodies showed a significant decrease in erlotinib-treated cells from 82.6% to 78% (P < 1E-4) for promoters and from 29.3% to 23.6% (P < 1E-4) for gene bodies, compared to control non-treated cells (Fig. 2b, Source Data file). Focusing on the 746 upregulated genes after erlotinib treatment (Fig. 1a-b and Supplementary Fig. 1c), we found that H3K27me3 levels were reduced in 63% (n = 470) of these genes including 24 TSG, whereas in only 256 genes (34%) including 9 TSG the H3K27me3 levels were increased (Fig. 2c; Supplementary Fig. 3c). Focusing on the H3K27me3 broad domains spanning these 470 upregulated genes (Fig. 2d; Source Data file) we detected significantly decreased H3K27me3 levels following erlotinib treatment at promoters (median = 4.64 RPMM; IQR = 3.96 RPMM; P = 8.87E-8) and gene bodies (median = 4.1 RPMM; IQR = 3.75 RPMM; P = 1.7E-2) as compared to the same promoters (median = 4.82 RPMM; IQR = 3.93 RPMM) and gene bodies (median = 4.56 RPMM; IQR = 3.88 RPMM) in Ctrl non-treated cells. Our results indicate that erlotinib treatment reduced genome-wide H3K27me3 levels including promoters and bodies of 470 genes that were upregulated following erlotinib treatment.

**Figure 2:**
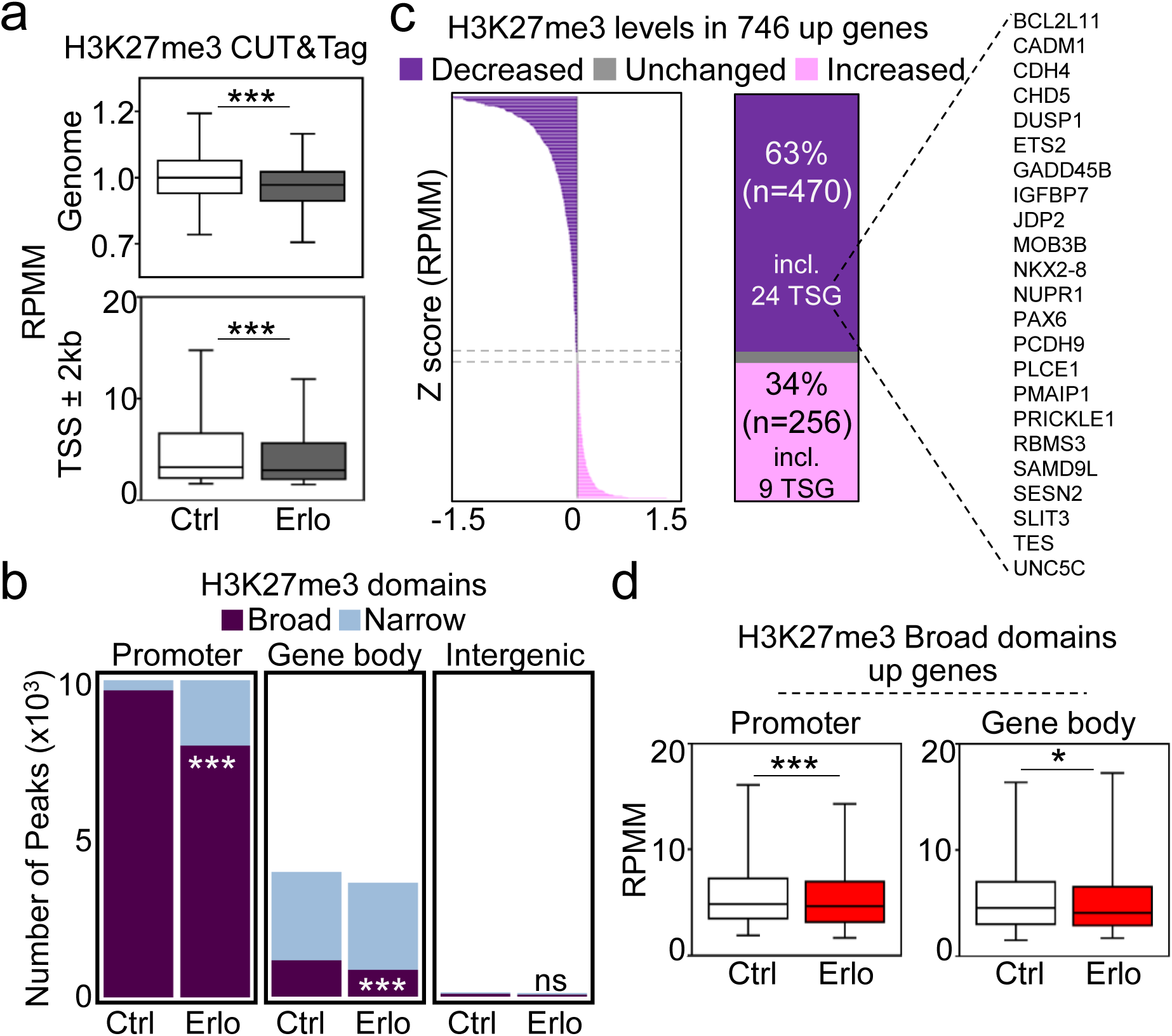
Gene activation induced by erlotinib correlates with reduced H3K27me3 broad domains. (**a**) Box plots showing the levels of H3K27me3 in control Ctrl or Erlo treated A549 cells genome-wide or at transcription start site (TSS -/+ 2 kb). Data were normalized using RPMM, read count per million mapped reads. (**b**) Bar plots displaying the broadness of H3K27me3 in different genomic regions at Promoters (Peaks -/+ 2 kb from TSS), Gene body (exon and intron regions outside the -/+ 2 kb TSS), and Intergenic (peaks not located in previous regions) in Ctrl or Erlo treated A549 cells. (**c**) Distribution of 746 upregulated genes based on H3K27me3 levels (63% decreased; n=470, including 24 TSG, 34% increased; n=256, including 9 TSG, and unchanged; n=20). Values, z-score of the normalized read counts as RPMM is obtained from annotatePeaks.pl from HOMER. (**d**) Box plots showing the levels of H3K27me3 in the significantly upregulated or downregulated genes that were separated into the indicated H3K27me3 domains (broad or narrow), and genomic regions (promoter or gene body) in Ctrl or Erlo treated A549 cells. All box plots display the median (middle line), 25th and 75th percentiles (box), and 5th and 95th percentiles (whiskers). Statistical significance is represented by asterisks: \*\*\**P* ≤ 0.001; \*\**P* ≤ 0.01; \**P* ≤ 0.05; ns, non-significant. *P*-values were calculated using a two-tailed t-test (box plots). See also Supplementary Fig. 3. Source data are provided as a Source Data file.

### Erlotinib activates enhancers and increases promoter-enhancer chromatin loops

As the majority of DEGs following erlotinib treatment in two different human pulmonary adenocarcinoma cell lines presented increased transcript levels (Fig. 1a-b, Supplementary Fig. 1c), we investigated the effect of erlotinib specifically in enhancers. We performed H3K27ac ChIP-seq in Ctrl non-treated or erlotinib treated A549 cells (Supplementary Fig. 4a) and analyzed the data using the rank-ordering of super-enhancers (ROSE) algorithm [68] to distinguish SE from TYE (Fig. 3a, left). We detected 1,649 SE in Ctrl non-treated A549 cells and this number significantly increased to 1,739 (P = 1.02E-4) following erlotinib treatment. Comparison of the genomic coordinates of the SE in Ctrl and erlotinib treated cells showed that only 701 SE were in common between both conditions (Fig. 3a, right). Furthermore, while we observed after erlotinib treatment significantly reduced H3K27ac levels in 948 SE that were specific to Ctrl cells (Fig. 3b, top; Source Data file), the levels of H3K27ac significantly increased in 1,038 SE that were specific for erlotinib treated cells (Fig. 3b, bottom; Source Data file). Genome-wide peak distribution analysis of the H3K27ac ChIP-seq data (Fig. 3c) showed that the increase of H3K27ac levels took place in intergenic regions, whereas the reduction of H3K27ac levels occurred at promoters and gene bodies, suggesting that most of the new SE after erlotinib treatment were in intergenic regions.

**Figure 3:**
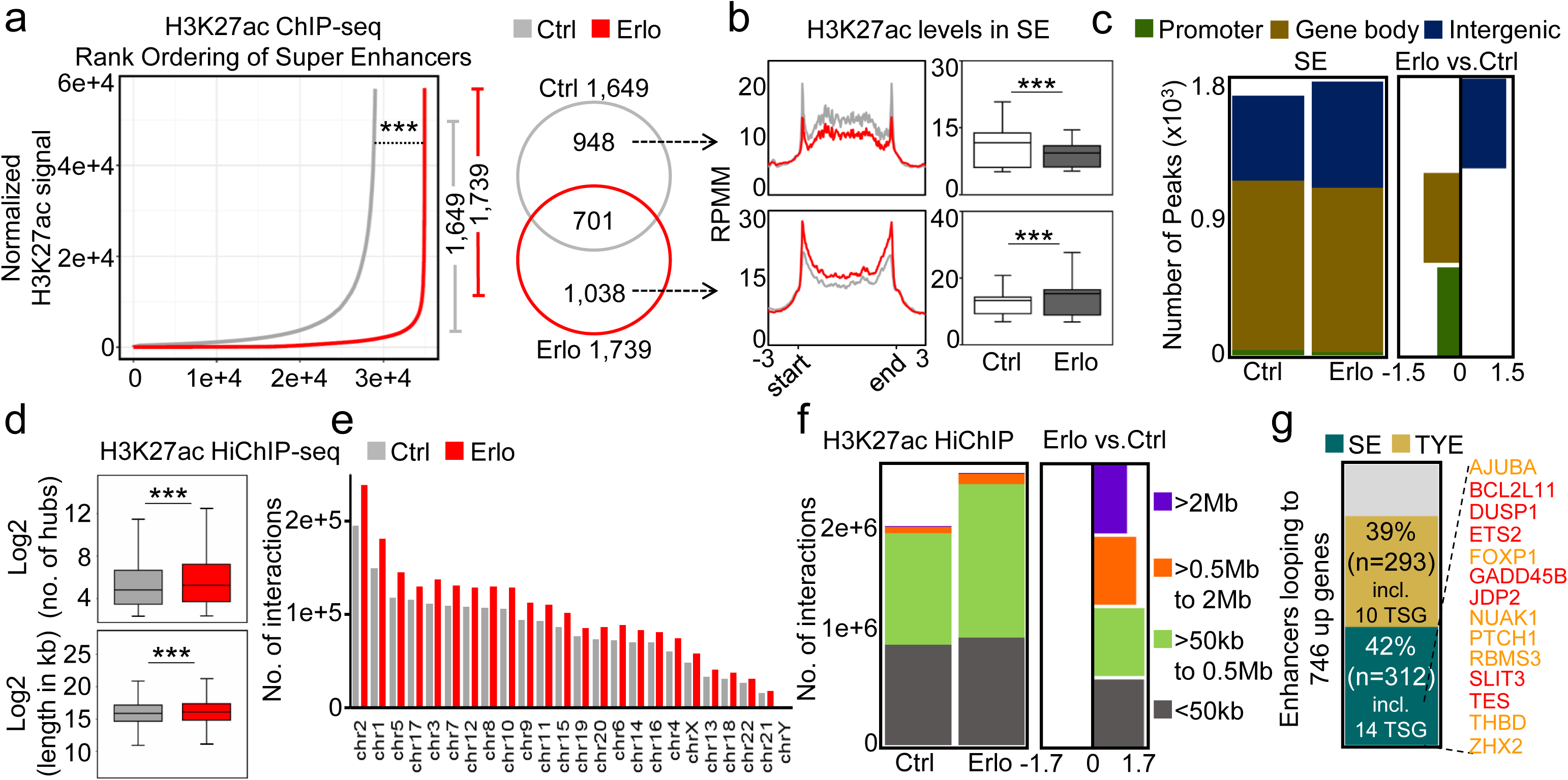
Erlotinib activates enhancers and increases promoter-enhancer chromatin loops. (**a**) Left, hockey stick plot after analysis using the ROSE algorithm and showing distribution of normalized H3K27ac ChIP-seq signal across typical enhancers (TYE) and super-enhancers (SE) in Ctrl (gray line) or Erlo (red line) treated A549 cells. Right, venn diagram showing the Ctrl-specific (n=948) and Erlo-specific (n=1,038) SE. (**b**) Aggregate and box plots after ChIP-seq in A549 cells showing the enrichment of H3K27ac and at Ctrl-specific or Erlo-specific SE. Data were normalized using RPMM, read count per million mapped reads. (**c**) Left, bar plots showing the distribution of H3K27ac peaks in different genomic regions at promoters, gene body, and intergenic regions. Right, shows H3K27ac enrichment in different genomic regions represented as ratios of peaks (Erlo versus Ctrl) in A549 cells. (**d**) Top, box plot showing the number of chromatin interaction hubs by H3K27ac-specific HiChIP-seq; bottom, box plot showing the size of the loops (length in kilobases, kb) in Ctrl or Erlo treated A549 cells. (**e**) Bar plot representing the total number of H3K27ac-specific chromatin interactions distributed across the chromosomes in the Ctrl and Erlo treated conditions. (**f**) Left, bar plots show the number of H3K27ac-specific chromatin interactions grouped based on the loop size as indicated (Mb; megabases, kb; kilobases). Right, squares represent the ratios of loop size (Erlo versus Ctrl). (**g**) Bar plot showing the distribution of enhancers looping to 746 upregulated genes (42% loop to SE; n=312, including 14 TSG, and 39% loop to TYE; n=293, including 10 TSG, unchanged). All box plots represent the median (middle line), 25th, 75th percentile (box), and 5th and 95th percentile (whiskers). In all plots asterisks represent *P*- values, \*\*\**P* ≤ 0.001; \*\**P* ≤ 0.01; \**P* ≤ 0.05; ns, non-significant. *P*-values were calculated after two-tailed t-test (box plots). See also Supplementary Fig. 4. Source data are provided as a Source Data file.

To determine the effect of erlotinib on 3D chromatin conformation we implemented a technique that combines in situ Hi-C library preparation with a chromatin immunoprecipitation (HiChIP, Fig. 3d-g, Supplementary Fig 4b-c, Source Data file). We used for this HiChIP-seq chromatin from Ctrl non-treated or erlotinib treated A549 cells and H3K27ac-specific antibodies to precipitate active enhancers that physically interact with promoters. We detected a significantly increased number of chromatin interaction hubs after erlotinib treatment (Fig. 3d, top) that were distributed across all chromosomes (Fig. 3e). Furthermore, the length of the chromatin loops also significantly increased after erlotinib treament (Fig. 3d, bottom; median = 16.11 kb; IQR = 2.53 kb; P = 2.22E-16) as compared to Ctrl non-treated cells (median = 15.88 kb; IQR = 2.5 kb). The increased number of chromatin loops following erlotinib treatment was more pronounced in chromatin loops exceeding 50 kb (Fig. 3f). Moreover, of the 746 upregulated genes (Fig. 1a-b, Supplementary Fig. 1c), we found 81% (n = 605) to physically interact through chromatin loops with active enhancers, including 42% (n = 312) with SE and 39% (n = 293) with TYE (Fig. 3g; Supplementary Fig. 4c; Source Data file).

### Erlotinib increases chromatin accessibility at Epromoters and super-enhancers bound by pioneer transcription factor FOXA2

Integrative analysis of the H3K4me3 ChIP-seq (Fig. 1e-i, Supplementary Fig. 2a-b), H3K27me3 CUT&Tag (Fig. 2a-d, Supplementary Fig. 3a-c) and H3K27ac HiChIP-seq (Fig. 3d-g, Supplementary Fig 4b-c) revealed that 214 genes among the 746 upregulated genes following erlotinib treatment (1) were located in H3K4me3 broad domains, with increased H3K4me3 levels, (2) have decreased H3K27me3 levels and (3) were physically interacting with active enhancers via chromatin loops (Fig. 4a), all these chromatin features likely contributing to their increased expression after erlotinib treament. Exploring the chromosome distribution of the 746 upregulated genes, we found that groups of these genes were in various chromosomes in same A compartments (Fig. 4b; Supplementary Fig. 5a) suggesting that genes in the same A compartment may be co-regulated. Visualization of the loci of various of these gene clusters using the integrative genome viewer (IGV) (Fig. 4c; Supplementary Fig. 5b), showed that genes in the same cluster were interacting through chromatin loops with each other and with enhancers. Moreover, the number and the length of the chromatin loops inside each cluster increased following erlotinib treatment, confirming our genome-wide results (Fig. 3d-g). Interestingly, analysis of publicly available ChIP-seq datasets [69,70] showed that the anchors of the chromatin loops were enriched with CTCF and components of the cohesin complex (RAD21 and SMC1A), suggesting their involvement in the regulation of chromatin changes induced by erlotinib.

**Figure 4:**
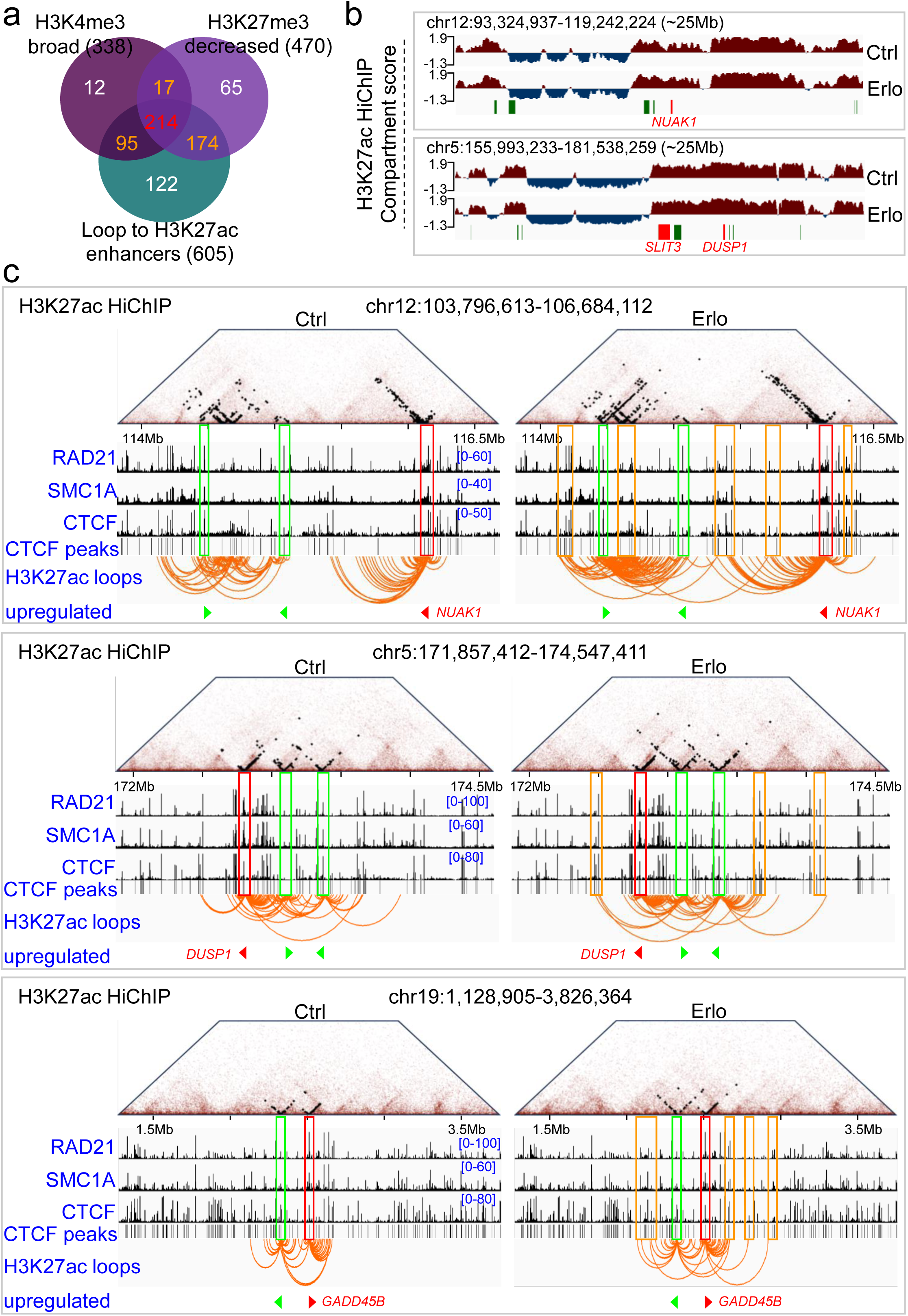
Erlotinib increases promoter-enhancer chromatin loops enriched with CTCF and the cohesin complex at their anchors. (**a**) Venn diagram showing the common upregulated genes (n=214) from the three groups, namely H3K4me3 broad (n=338); H3K27me3 decreased (n=470); and looping to enhancers (n=605). (**b**) Changes in chromatin compartmentalization are visualized across a ∼25Mb (Mb; megabases) window using compartment scores (y-axis) derived from H3K27ac-specific HiChIP experiments in the Ctrl and Erlo treated conditions. Upregulated genes in clusters are highlighted in green, and key tumor suppressor genes (TSG) are highlighted in red (*NUAK1*, *DUSP1*, and *SLIT3*). (**c**) Snapshots depict H3K27ac-specific chromatin contact matrices represented as pyramid plots comparing control (Ctrl, left) and erlotinib (Erlo, right) treated conditions at the top. Black dots within the pyramids represent the number of H3K27ac-specific interactions. Below the pyramid plots, ChIP-seq tracks illustrate factors of the cohesin complex (RAD21, SMC1A) and CTCF, along with CTCF peak annotations, and H3K27ac-specific chromatin loops. The visualization spans a ∼2.5Mb window surrounding the promoters of key TSG (red triangle) and upregulated gene in clusters (bright green triangles). Co-enrichment of RAD21, SMC1A, and CTCF at TSG promoter is highlighted in red boxes, while upregulated genes are marked with bright green boxes. Regions looping to the promoters are indicated by orange boxes, illustrating chromatin interactions. The IGV genome browser was used for visualization. See also Supplementary Fig. 5.

To gain further insight into these results, we analyzed sequencing data based on assay for transposase-accessible chromatin (ATAC-seq) in Ctrl or erlotinib treated pulmonary adenocarcinoma PC-9 cells [71] (Fig. 5a; Supplementary Fig. 6a-b; Source Data file). We found that erlotinib treatment significantly increased chromatin accessibility at TSS -/+1 kb of the 214 upregulated genes (Fig. 5a, top) and the 34 upregulated TSG (Fig. 5a, bottom). Motif search analysis of these TSS -/+1kb (Fig. 5b) revealed significant enrichment of nucleotide sequences containing binding motifs for the pioneer TF FOXA2 [7]. Confirming these results, FOAX2-specifc ChIP-seq (Fig. 5c; Supplementary Fig. 6c-d; Source Data file) showed significant enrichment of FOXA2 at the TSS -/+0.5kb of the 214 upregulated genes and the 34 upregulated tumor suppressor genes in a second pulmonary adenocarcinoma cell line (A549), whereas FOXA2 was not enriched at the same TSS in a human SCLC cell line (NCI-H889) supporting specificity of FOAX2 enrichment at TSS of the 746 upregulated genes with respect to the LC type (Supplementary Fig. 6d). Remarkably, similar results were obtained for the active, H3K27ac-labelled 1,038 SE (Fig. 3a-b) induced by erlotinib treatment (Fig. 5d-f). At these distal regulatory elements (SE), we observed increased chromatin accessibility following erlotinib treatment (Fig. 5d), significantly enriched nucleotide sequences containing FOXA2 binding motifs (Fig. 5e) that were confirmed by ChIP-seq showing FOXA2-enrichment at the active SE (Fig 5f). IGV genome browser snapshots focusing on the loci of the gene clusters previously shown in Fig. 4c and Supplementary Fig. 5b revealed increased chromatin accessibility following erlotinib treatment at the promoters of the upregulated genes, as well as at the enhancers that physically interacted with these promoters through chromatin loops (Fig. 6, top; Supplementary Fig. 6e, top). Interestingly, we observed inside each gene cluster that FOXA2 enrichment was higher in one of the promoters as compared to the promoters of other genes in the same cluster, specifically in pulmonary adenocarcinoma cells (Fig. 6, middle; Supplementary Fig. 6e, middle). Similarly, promoters with enhancer activity (Epromoters) that regulate transcription of various genes in a cluster are preferentially bound by a key TF [29,30]. Thus, we implemented a script to predict Epromoters [30] and found that those promoters with higher FOXA2 enrichment in each cluster were identified as Epromoters (Fig. 6, bottom; Supplementary Fig. 6e, bottom). Further, we analyzed an RNA-seq dataset in A549 cells containing long non-coding RNA molecules (50-2000 nucleotides) transcribed from enhancer regions (so called enhancer RNAs, eRNAs) [72,73], and found that eRNAs are transcribed from the predicted Epromoters (Fig. 6, bottom; Supplementary Fig. 6e, bottom), and with a lower frequency from the enhancers and other promoters in the same cluster. To confirm the results from the IGV genome browser snapshots of the loci of the gene clusters presented in Fig. 6 and Supplementary Fig. 6e, we predicted Epromoters [30] in the 746 upregulated genes and found 139 Epromoters with significantly increased chromatin accessibility after erlotinib treatment (Fig. 7a, Source Data file), significant enrichment of nucleotide sequences containing FOXA2 binding motifs (Fig. 7b, Source Data file) that was confirmed by ChIP-seq showing FOXA2-enrichment at Epromoters in pulmonary adenocarcinoma cells (A549) compared to the same loci in human SCLC cells (NCI-H889) supporting specificity of FOAX2 enrichment at these 139 Epromoters with respect to the lung cancer type (Fig 7c, Source Data file). Moreover, RNA-seq-based transcriptome analysis in cells that were transfected with Ctrl or *FOXA2*-specific small interfering RNAs (siRNA, *siCtrl*, *siFOXA2*; Fig. 7d, Supplementary Fig. 6f; Source Data file) showed a significant decrease in the expression levels of the 746 upregulated transcripts, including the 214 upregulated genes identified in Fig. 4a and the 34 TSG, after *FOXA2*-specific loss-of-function (LOF; Fig 7d) demonstrating the requirement of *FOXA2* for the basal transcriptional levels of these genes. Our results indicate that the genes in the same clusters are (1) co-activated after erlotinib treatment through increased chromatin accessibility at TSS, (2) binding of FOXA2 preferentially to an Epromoter in each cluster, (3) physical interaction with active SE via chromatin loops that are stabilized by CTCF and the cohesin complex.

**Figure 5:**
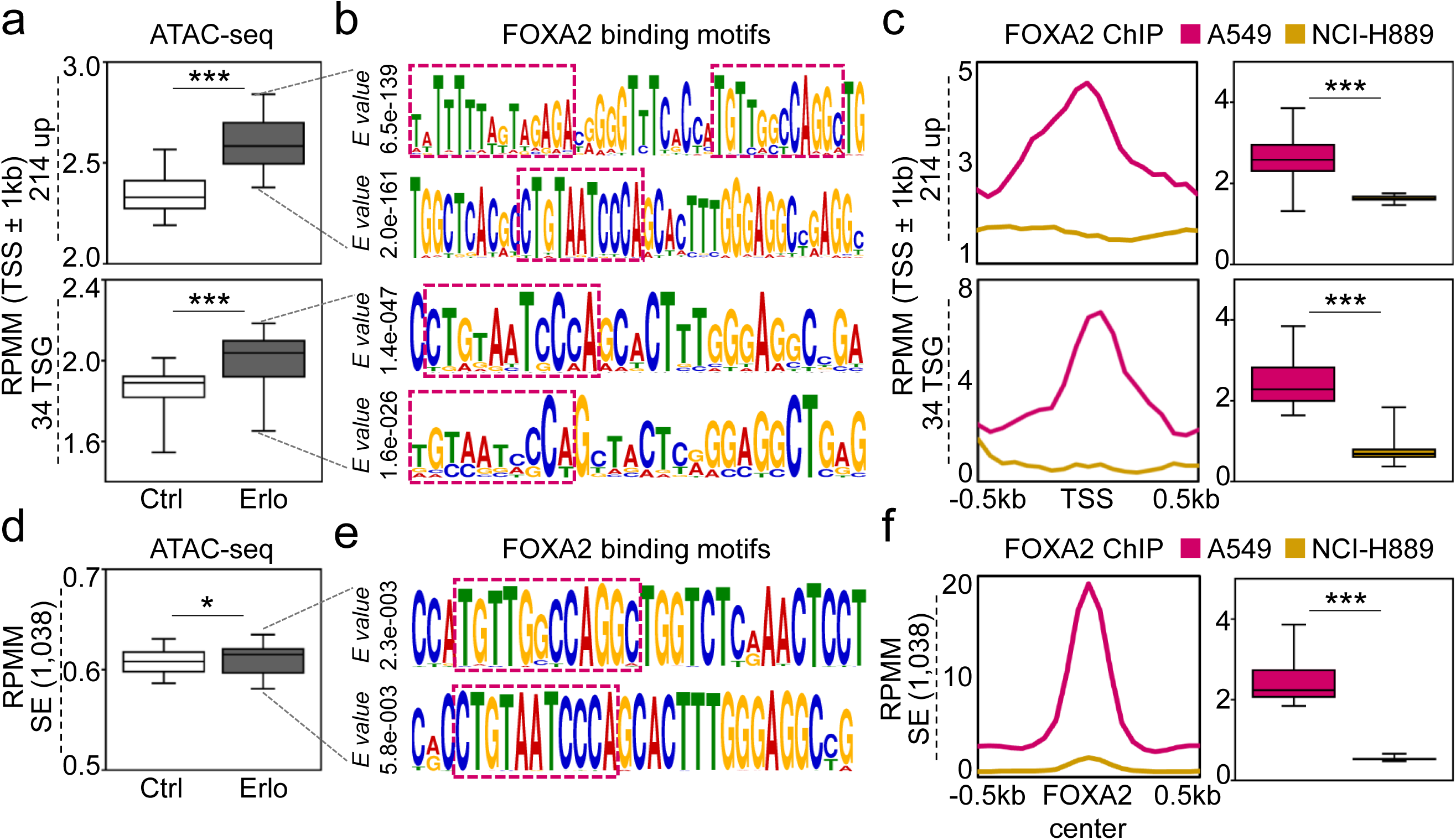
Erlotinib increases chromatin accessibility at TSS and super-enhancers bound by pioneer transcription factor FOXA2. (**a**) Box plots from ATAC-seq showing the accessibility at the transcription start site (TSS -/+ 1 kb) of 214 upregulated genes (top) or 34 tumor suppressor genes (TSG; bottom) in Ctrl or Erlo treated PC-9 cells. Data were normalized using RPMM, read count per million mapped reads. (**b**) Motif analysis reveals significant enrichment of nucleotide motifs from 214 upregulated genes and 34 TSG that show FOXA2 binding sites (indicated by magenta dotted boxes) as predicted using the JASPAR database. *E value* represents the statistical significance of the motif. (**c**) Aggregate and box plots showing the FOXA2 ChIP-seq enrichment at transcription start site (TSS -/+ 1 kb) of 214 upregulated genes or 34 TSG in A549 and NCI-H889 cells. (**d**) Similar to **a**, box plots from ATAC-seq showing accessibility at loci of super enhancers (SEs, n=1,038). (**e**) Similar to **b**, motif analysis highlights significant enrichment of nucleotide motifs at SE loci, revealing FOXA2 binding sites. (**f**) Similar to **c**, aggregate and box plots showing FOXA2 enrichment at SE loci. All box plots display the median (middle line), 25th and 75th percentiles (box), and 5th and 95th percentiles (whiskers). Statistical significance is represented by asterisks: \*\*\**P* ≤ 0.001; \*\**P* ≤ 0.01; \**P* ≤ 0.05; ns, non-significant. *P*-values were calculated using a two-tailed t-test (box plots). See also Supplementary Fig. 6. Source data are provided as a Source Data file.

**Figure 6:**
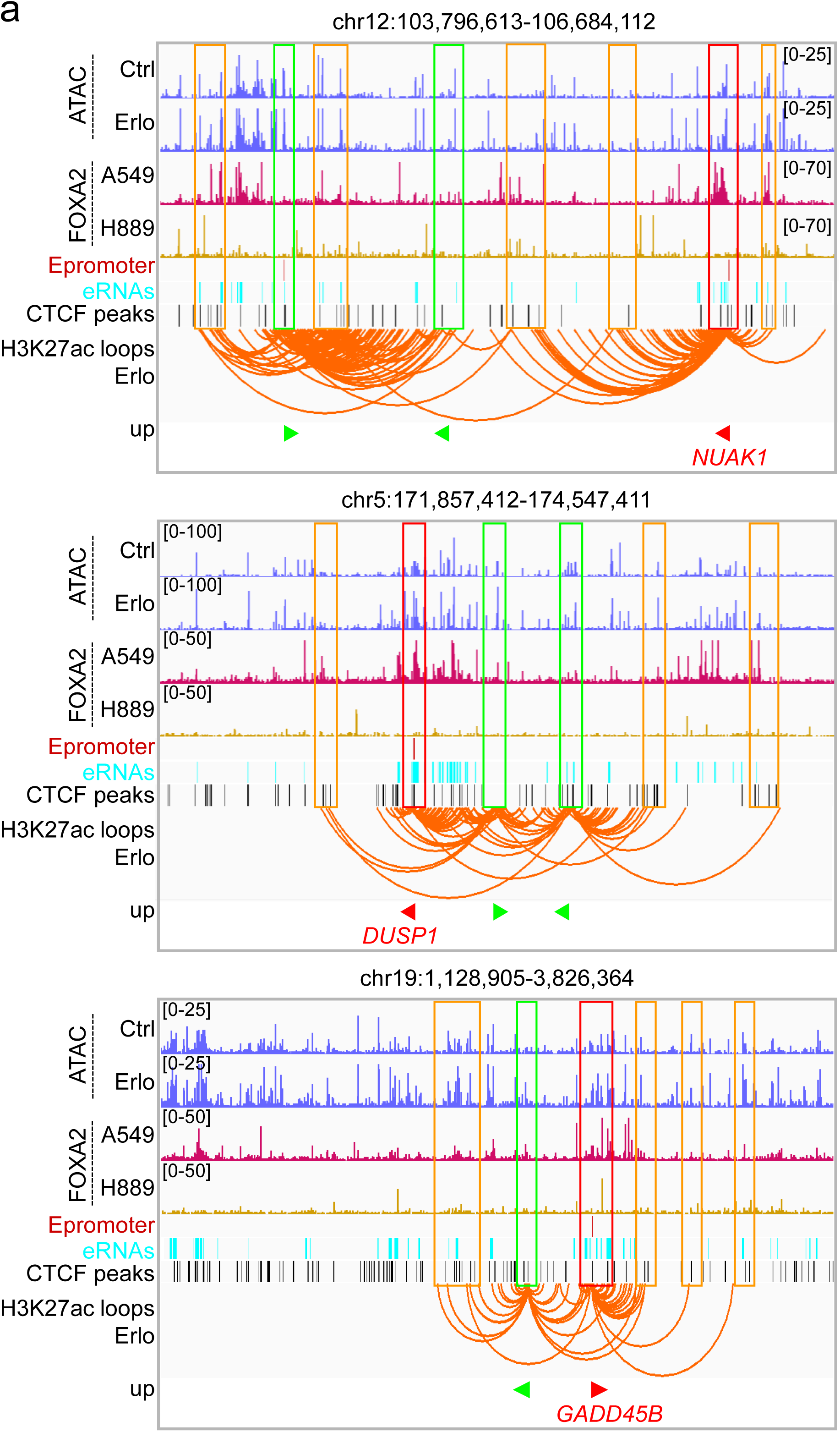
Erlotinib increases chromatin accessibility at Epromoters and enhancers bound by pioneer transcription factor FOXA2. (**a**) Genome browser snapshots of selected tumor suppressor genes (TSG) loci showing chromatin accessibility by ATAC-seq (blue) in Ctrl and Erlo treated PC-9 cells, FOXA2 ChIP-seq in A549 (magenta) and NCI-H889 cells (gold). The tracks below depict Epromoters (red), eRNAs (arctic blue), CTCF peaks (black), and H3K27ac-specific chromatin loops in Erlo treated A549 cells. The visualization spans a ∼2.5Mb window surrounding the promoters of key TSG (red triangle) and upregulated gene in clusters (bright green triangles). Co-enrichment of FOXA2 with ATAC-seq, Epromoter, eRNAs, and CTCF peaks at TSG promoter is highlighted in red boxes, while upregulated genes are marked with bright green boxes. Regions looping to the promoters are indicated by orange boxes, illustrating chromatin interactions. The IGV genome browser was used for visualization. See also Supplementary Fig. 6.

**Figure 7:**
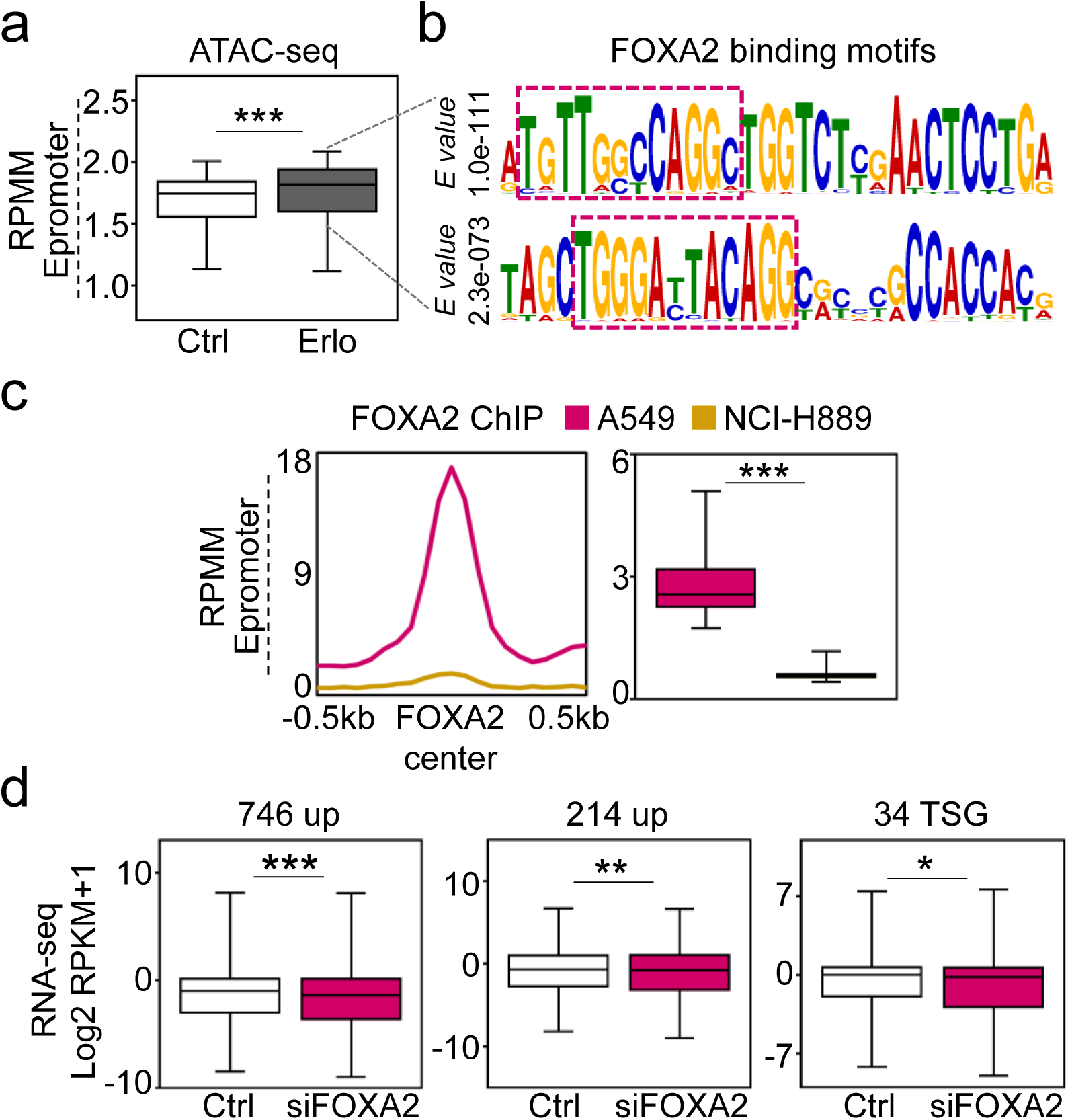
Erlotinib increases chromatin accessibility at Epromoters bound by pioneer transcription factor FOXA2. (**a**) Box plots from ATAC-seq showing the accessibility at Epromoters (n=139) in Ctrl or Erlo treated PC-9 cells. Data were normalized using RPMM, read count per million mapped reads. (**b**) Motif analysis reveals significant enrichment of nucleotide motifs Epromoters that show FOXA2 binding sites (indicated by magenta dotted boxes) as predicted using the JASPAR database. *E value* represents the statistical significance of the motif. (**c**) Aggregate and box plots showing the FOXA2 ChIP-seq enrichment at Epromoters in A549 and NCI-H889 cells. (**d**) Box plots of RNA-seq-based expression analysis of upregulated genes (left; n=746, middle; n=214) and tumor suppressor genes (TSG; n=34) in the control (Ctrl) or siRNA construct specific for FOXA2 (siFOXA2). All box plots display the median (middle line), 25th and 75th percentiles (box), and 5th and 95th percentiles (whiskers). Statistical significance is represented by asterisks: \*\*\**P* ≤ 0.001; \*\**P* ≤ 0.01; \**P* ≤ 0.05; ns, non-significant. *P*-values were calculated using a two-tailed t-test (box plots). See also Supplementary Fig. 6. Source data are provided as a Source Data file.

### Gene expression signature induced by erlotinib can be used for diagnosis and prognosis of LUAD patients

Extending our findings to translational applicability, we performed principal component analysis (PCA) of RNA-seq data from Ctrl donors and NSCLC patients deposited at TCGA restricted to 746 upregulated genes following erlotinib treatment (Fig. 1a-b; Supplementary Fig. 1c). This analysis not only allowed us to differentiate between Ctrl donors and NSCLC patients, but also to discriminate between the NSCLC subtypes LUAD and LUSC (Fig. 8a, left). Similar results were obtained after PCA of RNA-seq data deposited at TCGA using the expression of 214 upregulated genes identified in Fig. 4a (Fig. 8a, right). Further, genome-wide association studies (GWAS) of cancer genomics datasets revealed 960 single nucleotide polymorphisms (SNPs) in the 34 TSG included in the 746 upregulated genes (Fig. 8b). Interestingly, 26% (n = 246) of these SNPs were found in LC genomics datasets, from which LUAD was the most frequent (16%; n = 152) followed by LUSC (5%; n = 49) (Fig. 8c). The most frequent mutation type in the LC-specific SNPs was missense mutations (75%; n = 185) (Fig. 8d). Similarly, GWAS of cancer genomics datasets focusing on the 1,038 SE induced after erlotinib treatment revealed 164 SNPs, from which 22% (n = 36) were found in LC genomics datasets (Fig. 8e). Interestingly, 44% (n = 16) of these SNPs were found in LUAD genomics datasets, followed by LUSC data sets (19%; n = 7) (Fig. 8f). Survival analysis of the 1,411 LC patients from the Kaplan-Meier plotter [74] (Fig 8g, left; Source Data file) showed a significantly longer survival of patients with increased levels of the 34 TSG included in the 746 upregulated genes after erlotinib treatment (n = 704; median survival = 99.4 months; hazard ratio = 0.62; P = 2.5E-10; Cox regression) as compared to patients with low expression levels (n = 707; median survival = 52 months). Remarkably, the positive effect of the increased levels of the 34 TSG on the longer survival of patients was specific to LUAD patients (Fig. 8e, middle; Source Data file), since it was not observed in LUSC patients (Fig. 8e, right; Source Data file). Similar results were obtained when the *FOXA2* expression levels were considered for survival analysis in the 2,166 LC patients from the Kaplan-Meier plotter (Supplementary Fig. 7a-c; Source Data file). These findings support the clinical relevance of the 746 genes upregulated following erlotinib treatment identified here (Fig. 1a-b; Supplementary Fig. 1c) as gene expression signature not only for molecular differentiation of NSCLC patients in LUAD and LUSC, but also for prognosis prediction of LUAD patients, which might help to develop patient-tailored therapies.

**Figure 8:**
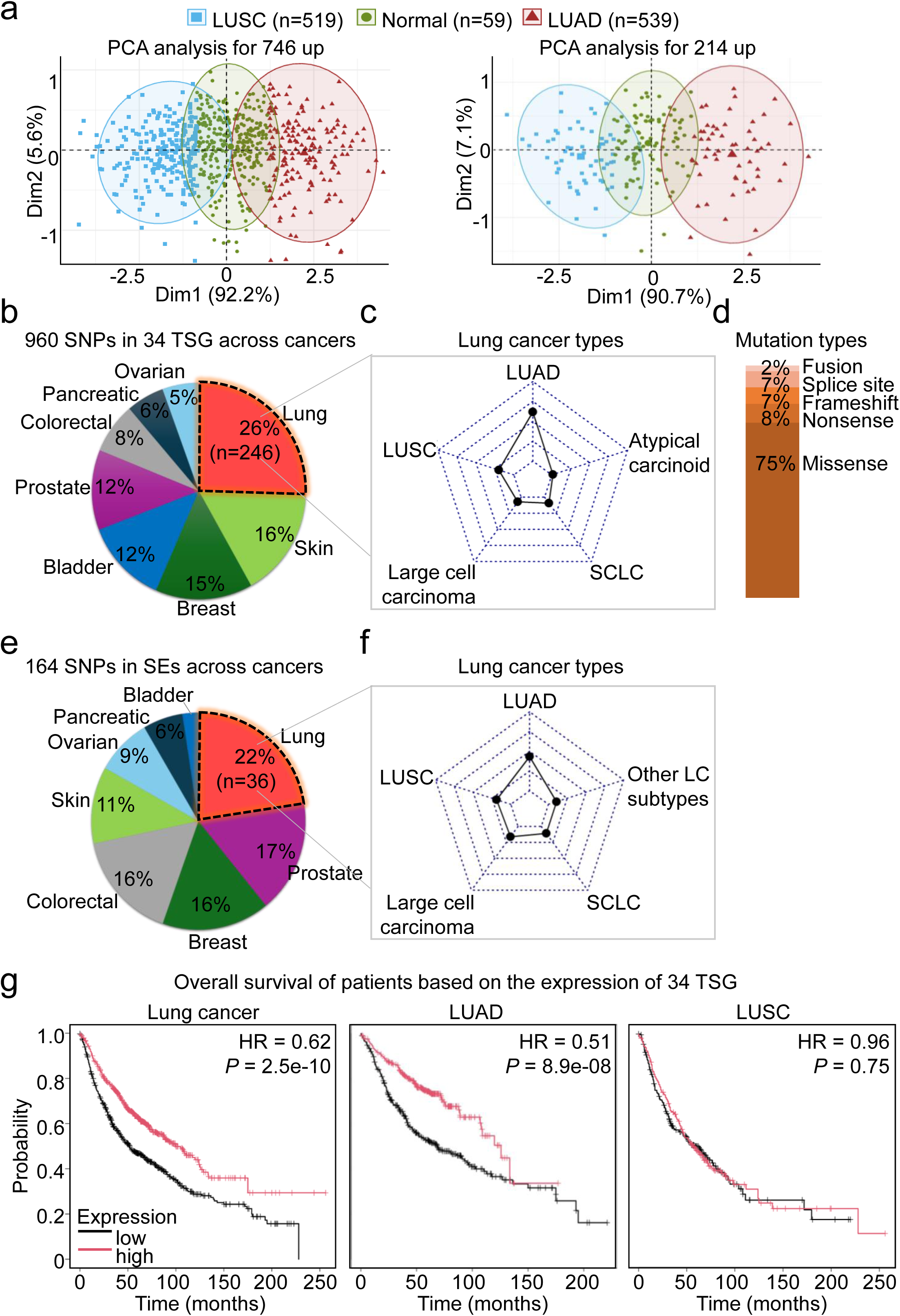
Gene expression signature induced by erlotinib can be used for diagnosis and prognosis of LUAD patients. (**a**) Principal Component Analysis (PCA) plots from highlighting the differences in gene expression patterns among three sample groups from TCGA: Lung Squamous Cell Carcinoma patients (LUSC; n=519, blue squares), Normal lung samples (Normal; n=59, green circles), and Lung Adenocarcinoma patients (LUAD; n=539, brown triangles). The left PCA plot illustrates data distribution along Dim1 (92.2% variance) and Dim2 (5.6% variance) for the 746 upregulated genes, while the right PCA plot shows distribution along Dim1 (90.7% variance) and Dim2 (7.1% variance) for the 214 upregulated genes. (**b, c, d**) Pie chart showing the distribution of 960 single nucleotide polymorphisms (SNPs) across 34 tumor suppressor genes (TSGs) in various cancers. The segments represent different cancers, with lung cancer showing the highest proportion of SNPs (26%, n=246), followed by other cancer types. In **c**, the radar plot visualizes the 246 SNPs across different lung cancer types, including LUAD (n=152), LUSC (n=49), Large cell carcinoma (n=23), SCLC (Small Cell Lung Cancer, n=18), and atypical carcinoid (n=4). In **d**, bar plot illustrates the types of mutations in lung cancer, the mutation categories include fusion, splice site, frameshift, nonsense, and missense as indicated. (**e, f**) Pie chart showing the distribution of 164 SNPs across super enhancers (SEs; n=1,038) in various cancers. The segments represent different cancers, with lung cancer showing the highest proportion of SNPs (22%, n=36), followed by other cancer types. In **f**, the radar plot visualizes the 36 SNPs across different lung cancer types, including LUAD (n=16), LUSC (n=7), Large cell carcinoma (n=6), SCLC (n=4), and other LC types (n=3). (**g**) Kaplan-Meier survival curves illustrating overall survival of patients based on the expression of 34 TSGs across different lung cancer types. Survival probabilities are represented for all lung cancer patients (left), LUAD patients (middle), and LUSC patients (right). Patients with high TSG expression (red curves) show distinct survival patterns compared to those with low expression (black curves). Lung cancer (HR = 0.62, *P* = 2.5e-10, median survival in months: high expression vs. low expression is 99.43 vs. 52), LUAD (HR = 0.51, *P* = 8.9e-08, median survival in months: high expression vs. low expression is 125.7 vs. 69), and LUSC (HR = 0.96, *P* = 0.75, median survival in months: high expression vs. low expression is 57 vs. 58). HR, hazard ratio. See also Supplementary Fig. 6. Source data are provided as a Source Data file.

## DISCUSSION

Geusz and colleagues demonstrated a dual role for FOXA pioneer TFs in endodermal organ development [26]. First, in endodermal organ precursor cells, FOXA pioneer TFs bind to primed enhancers, which are enriched for strong FOXA binding motifs, as well as for binding motifs of cell-lineage-specific TFs mediating response to determined signaling pathways, such as EGF signaling. The binding of FOXA pioneer TFs to these primed enhancers induced chromatin rearrangements resulting in a more accessible chromatin state at genomic elements regulating organ cell type-specific gene expression. Second, this accessible chromatin state allows cell-lineage-specific TFs to bind these regulatory elements and modulate gene expression facilitating signal-dependent lineage initiation and enforcing organ cell type-specific gene expression. Our results support the hypothesis that FOXA pioneer TFs play a similar dual role in NSCLC and regulate chromatin rearrangements that allow binding of cell-lineage-specific TFs mediating response to EGF signaling to induce cancer-related gene expression signatures. Supporting this hypothesis, we showed that blocking EGF signaling in NSCLC cells by EGFR-TKIs induce changes in their 3D genome, including increased H3K4me3 broad domains, reduced H3K27me3 levels and increased chromatin loops mediating promoter-SE interactions. These chromatin rearrangements together lead to increased expression of 746 genes including 34 TSG that may explain the therapeutic effects of the EGFR-TKIs in LUAD. The observed changes in chromatin of NSCLC cells after EGFR-TKI treatment may be related to the pioneer TF FOXA2, which is enriched at the TSS of the 34 increased TSG (Fig. 5c), active SE (Fig. 5f) and Epromoters regulating cluster of upregulated genes after erlotinib treatment (Fig. 7c). Interestingly, inhibition of EGFR by another TKI (afatinib) alone or in combination with gemcitabine (nucleoside analog used as chemotherapy medication) [75] led to eradication of cancer stem cells and reduced pancreatic cancer metastasis [77]. Since FOXA pioneer TFs are required for embryonic development of organs that arise from the endoderm [32–36], it will be interesting to determine whether EGFR-TKIs will induce similar changes in the 3D genome of pancreatic cancer cells, or of cancer cells in other organs of endodermal origine, similar to the 3D genome rearrangements reported here for lung cancer cells. The implications of embryonic master regulators in promoting tumorigenesis can be exploited clinically as new therapeutic targets to specifically ablate cancer cells. Following this line of ideas, specific silencing of the pioneer factor FOXA1 in cancer cells reduced cancer hallmarks [76].

The *EGFR* mutations ΔEx19 and L858R are the most common genomic drivers of NSCLC that are targetable with first-generation EGFR-TKIs, such as erlotinib and gefitinib. However, most of the NSCLC patients treated with these EGFR-TKIs develop resistance to the treatment. Over 50% of the patients with acquired EGFR-TKI resistance harbor a secondary point mutation in the EGFR kinase domain that substitutes threonine 790 by methionine (T790M) producing a drug-resistant EGFR variant [77]. The need to overcome this resistance mechanism led to the development of third-generation EGFR-TKIs, of which osimertinib is currently the only one with regulatory approval. Osimertinib is an EGFR (T790M) mutant selective EGFR-TKI that induces a similar gene expression signature as erlotinib and gefitinib (Supplementary Fig. 1c). It is reasonable to assume that osimertinib will induce similar effects on the 3D genome as the ones described here for erlotinib and gefitinib. Thus, our results showing the effect of EGFR-TKIs on different chromatin features provide the molecular basis for designing more specific therapies targeting H3K4me3 broad domains, SE or Epromoters that will allow to induce the expression of TSG. Building on this perspective, broad H3K4me3 domains have been linked to increased transcription elongation and enhancer activity, collectively leading to the high expression of the TSG *TP53* and *PTEN* [11]. Further, SE were found at key oncogenic drivers and other genes that are important in tumor pathogenesis [24]. Interestingly, treatment of multiple myeloma cells with the BET-bromodomain inhibitor JQ1 led to loss of the transcriptional coactivator BRD4 at SE and consequent transcription elongation defects, that preferentially impacted oncogenes, including the *MYC* oncogene [78]. Another study by Lewis and colleagues showed that deletion of a SE or specific degradation of its eRNA reduced the expression of the SE-controlled *PODXL* gene, thereby suppressing cell proliferation, migration, tumor growth, and metastasis in mouse xenograft models of triple-negative breast cancer [79]. Moreover, Wang and colleagues found by integrating GWAS, expression Quantitative Trait Loci (eQTLs) and 3D chromatin interactions that genetic variations at Epromoters are associated with various cancer types [80]. It will be the scope of our future work to determine the clinical relevance of targeting H3K4me3 broad domains, SE or Epromoters using CRISPR–Cas9 gene-editing technology [81] either to reduce oncogene expression, or to induce TSG expression as therapeutic strategy for NSCLC alone or as combination treatment to improve the efficacy of treatment with EGFR TKIs.

## METHODS

### Cell culture

Human NSCLC epithelial cells A549 (ATCC CCL-185) were cultured in complete RPMI 1640 medium supplemented with 10% FBS, 1% penicillin-streptomycin, and 2mM L-glutamine. Cells were maintained at 37 °C, 5% CO_2_ in a humidified incubator. During subculturing, cells were washed with 1x PBS trypsinized with 0.25% (w/v) Trypsin and subcultured at the ratio of 1:5 to 1:10. The cell lines used in this paper are mycoplasma free. They were regularly tested for mycoplasma contamination. In addition, they are not listed in the database of commonly misidentified cell lines maintained by ICLAC.

### Erlotinib treatment

A549 cells were cultured RPMI 1640 medium with standard 100 mm cell culture dishes. Erlotinib hydrochloride (Sigma-Aldrich #SML2156) was dissolved in DMSO (Sigma-Aldrich #34869) according to the manufacturer’s instructions and used at a final concentration of 5 µM. The cells were treated with 5 µM erlotinib (Erlo) or DMSO as a control (Ctrl) for 48 h. After 24 h of treatment, the medium was replenished with fresh erlotinib to maintain consistent treatment conditions.

### Total RNA isolation and RNA sequencing data analysis

Total RNA from A549 cells were isolated using the RNeasy Mini kit (Qiagen), quantified using a Nanodrop Spectrophotometer (ThermoFisher Scientific), and was subjected to total RNA sequencing. RNA sequencing data for this paper were generated as previously described [16,82]. Total RNA and library integrity were verified on LabChip GX Touch 24 (Perkin Elmer). Sequencing was performed on the NextSeq500 instrument (Illumina) using V2 chemistry with paired end setup. Raw reads were visualized by FastQC (https://www.bioinformatics.babraham.ac.uk/projects/fastqc/) to determine the quality of the sequencing. Trimming was performed using trimmomatic (version 0.39) [83] with the following parameters LEADING:3 TRAILING:3 SLIDINGWINDOW:4:15 HEADCROP:5 MINLEN:15. High quality reads were mapped to the human genome (hg38) using Bowtie2 (version 2.4.4) [84]. SAM files were sorted and converted to BAM files using Samtools (version 1.13) with the command samtools view -Sb -u (https://www.htslib.org/doc/samtools-view.html). BAM files were converted to BED files using bedtools (https://bedtools.readthedocs.io/en/latest/content/tools/bamtobed.html). After mapping, tag directories were obtained with makeTagDirectory from HOMER (version 4.11.1) (http://homer.ucsd.edu/homer/ngs/tagDir.html) using default settings [85]. Samples were then quantified by using analyzeRepeats.pl (http://homer.ucsd.edu/homer/ngs/analyzeRNA.html) from HOMER with the parameters (analyzeRepeats.pl rna hg38 -count genes -d Tag -rpkm; reads per kilobase per millions mapped). To avoid division through zero, those reads with zero RPKM were set to 0.001. Upregulated genes after erlotinib treatment defined for those genes with a log2FC (Erlotinib/Control) ≥ 0.5 and downregulated those genes with a log2FC (Erlotinib/Control) ≤ 0.5. Volcano and all box plots were generated using GraphPad Prism 8 software.

### Chromatin immunoprecipitation (ChIP)

ChIP analysis was performed as described earlier with minor adaptations [21]. Briefly, cells were cross-linked with 1% methanol-free formaldehyde (ThermoFisher Scientific) lysed, and sonicated with Diagenode Bioruptor to an average DNA length of 300-600 bp. After centrifugation, the soluble chromatin was immunoprecipitated with 2 µg of antibodies specific for H3K4me3 (Abcam, #ab8580), H3K27ac (Abcam, #ab4729), and IgG (Santa Cruz, #sc-2027). Immunoprecipitated chromatin was purified using the QIAquick PCR purification kit (Qiagen) and subjected to next-generation sequencing. TruSeq DNA library preparation kit (Illumina) was used to generate the ChIP libraries and sequenced using Illumina HiSeq 2500 system.

### Cleavage under targets and tagmentation (CUT&Tag)

CUT&Tag experiments were performed as described previously [86], using the antibody specific for H3K27me3 (Cell Signaling Technology, #9733). Briefly, 1 × 10^6^ cells were harvested, washed with wash buffer (20 mM HEPES pH 7.5, 150 mM NaCl, 0.5 mM spermidine), and immobilized to concanavalin A coated beads with incubation at room temperature for 10 min. The bead-bound cells were incubated in 200 µl of primary antibody buffer (wash buffer with 1% BSA, 2 mM EDTA, and 0.05% digitonin for gentle permeabilization of the plasma and nuclear membrane) with 1:100 primary antibody dilution at 4 °C by rotating overnight. The next day, the primary antibody buffer was removed and cells were washed with 800 µl of dig-wash buffer (wash buffer with 1% BSA and 0.05% digitonin) three times to remove unbound antibodies. The cells were then incubated with guinea pig anti-rabbit antibody (Novus Biologicals, NBP1-72763) in 200 µl of dig-wash buffer at room temperature for 1 h with slow rotation. After a brief wash with dig-wash buffer as above, cells were resuspended in 200 µl of dig-300 buffer (20 mM HEPES pH 7.5, 300 mM NaCl and 0.5 mM spermidine, 1% BSA and 0.01% digitonin) with 1:200 dilution of CUT&Tag-IT^®^ Assembled pA-Tn5 Transposomes (Active Motif, #53164) and incubated at room temperature for 1 h with slow rotation. pA-Tn5-bound cells were washed with 800 µl of dig-300 buffer three times, followed by tagmentation in 200 µl of tagmentation buffer (dig-300 buffer with 10 mM MgCl_2_) at 37 °C for 1 h. To stop tagmentation, and solubilize DNA fragments, add 10 µL 0.5M EDTA, 3 µL 10% SDS and 2.5 µL 20 mg/mL Proteinase K were added to each sample and mixed by full-speed quick vortexing, and further incubated at 63 °C for another 1 h to digest protein and to reverse cross-link DNA. Genomic DNA was extracted and purified QIAquick PCR purification kit (Qiagen) and subjected to library preparation as indicated (https:/www.protocols.io/view/bench-top-cut-and-tag) with minor adaptations. The libraries were size selected by SPRI-bead-based approach after the final PCR with 18 cycles. In detail, samples were 1st cleaned up by 1x bead:DNA ratio to eliminate residuals from PCR reaction, followed by a 2-sided-bead cleanup step with an initial 0.6x bead:DNA ratio to exclude larger fragments. The supernatant was transferred to a new tube and incubated with additional beads in 0.2x bead:DNA ratio to eliminate smaller fragments, like adapter and primer dimers. Bound DNA samples were washed with 80% ethanol, dried, and resuspended in TE buffer. Library integrity was verified with 2100 Bioanalyzer system (Agilent Technologies). TruSeq DNA library preparation kit (Illumina) was used to generate the CUT&Tag libraries and sequenced (paired-end) using Illumina HiSeq 2500 system.

### Meta-analysis of NGS data (ChIP-seq, CUT&Tag, and ATAC-seq)

Raw reads of samples from ChIP-seq, CUT&Tag, and ATAC-seq were visualized by FastQC (https://www.bioinformatics.babraham.ac.uk/projects/fastqc/) to determine the quality of the sequencing. Trimming was performed using trimmomatic (version 0.39) [83] with the following parameters LEADING:3 TRAILING:3 SLIDINGWINDOW:4:15 HEADCROP:5 MINLEN:15. High quality reads were mapped to the human genome (hg38) using Bowtie2 (version 2.4.4) [84]. For the CUT&Tag samples, the trimmed reads were mapped with bowtie2 settings (--local --very-sensitive --no-mixed --no-discordant --phred33 -I 10 -X 700). SAM files were sorted and converted to BAM files using Samtools (version 1.13) with the command samtools view -Sb -u (https://www.htslib.org/doc/samtools-view.html). Further, PCR duplicated reads were removed using picard tools (version 3.3.0) (https://broadinstitute.github.io/picard/command-line-overview.html#MarkDuplicates).

Duplicate-removed BAM files were converted to BED files using bedtools (https://bedtools.readthedocs.io/en/latest/content/tools/bamtobed.html). The BED files were further used to create tag directories using the function makeTagDirectory (default settings) from HOMER (version 4.11.1) (http://homer.ucsd.edu/homer/ngs/tagDir.html). Reads Per kilobase per million mapped reads (RPKM) normalized coverage tracks were generated using deepTools (version 3.5.6) [87] with the bamCoverage function. Bedtools multicov (https://bedtools.readthedocs.io/en/latest/content/tools/multicov.html) was used to quantify the signal from BAM files that overlap with intervals specified in a BED file. Peak calling for H3K4me3 was performed as described previously [12] using MACS2 (version 2.2.9) with settings (callpeak -g hs -q 0.05 --broad --extsize 1000 --keep-dup 1 --nomodel) [88]. To define H3K4me3 broad domains, summary statistics of peak sizes were initially analyzed. Peaks larger than 2.7 kilobases (kb) were classified as broad domains, while those smaller than 1.9 kb were categorized as narrow domains. Peak calling for H3K27ac was performed with the settings (callpeak -g hs -q 0.05 --broad –extsize 200 --keep-dup 1 --nomodel). Blacklisted regions were subtracted from the peaks using bedtools intersect -v option (https://bedtools.readthedocs.io/en/latest/content/tools/intersect.html). The peaks were then annotated to the genomic regions of human genome with the function annotatePeaks.pl (http://homer.ucsd.edu/homer/ngs/annotation.html) from HOMER. Aggregate or enrichment plots and heatmaps to quantify ChIP-seq, CUT&Tag, ATAC-seq signals normalized by read count per million mapped reads (RPMM) or RPKM were generated from deepTools using their computeMatrix command followed by plotProfile or plotHeatmap functions or using the function annotatePeaks.pl from HOMER (http://homer.ucsd.edu/homer/ngs/quantification.html) with the parameters (annotatePeaks.pl hg38 -size 2000 -hist 50 -d Tag -norm). The enrichment was quantified around the promoters at TSS ± 1kb/TSS ± 2kb, genome wide or at defined coordinates as indicated in the figures. Quantified enrichment scores were used to generate box plots.

For the visualization of tracks, including BigWig or BedGraph tracks, peaks, and BED files, as well as for generating snapshots, the Integrative Genomics Viewer (IGV) was used [89].

### Definition of Super-Enhancers and Epromoters

To identify super-enhancers, H3K27ac ChIP-seq signal and peaks were used as inputs for the program Rank Ordering of Super-Enhancer (ROSE package) with default settings [68,78]. To define Ctrl-specific and Erlo-specific super-enhancers, the respective peaks obtained using MACS2 were used as inputs. The Ctrl peaks were called by using the Erlo BAM file as the background, while the Erlo peaks were called by using the Ctrl BAM file as the background. Epromoters were defined using the script described earlier, with minor adaptations [30]. H3K27ac ChIP peaks and promoters (1kb upstream from TSS) were used as inputs for the script, and the Epromoters were crossed with FOXA2 peaks using bedtools intersect.

### Definition of H3K27me3 broad domains

Peak calling for H3K27me3 broad domains was performed with the settings (callpeak -g hs -q 0.05 --broad --extsize 1000 --keep-dup 1 --nomodel). Then, peaks were then merged to a maximum distance of 4 kb using bedtools with the option bedtools merge -d 4000 (https://bedtools.readthedocs.io/en/latest/content/tools/merge.html). To define H3K27me3 broad domains, summary statistics of peak sizes were initially analyzed. Peaks larger than 1.9 kb were classified as broad domains, while those smaller than 1.5 kb were categorized as narrow domains.

### Chromatin conformation capture by in situ Hi-C followed by chromatin immunoprecipitation (HiChIP)

HiChIP experiments were performed as previously described [90] using antibodies specific for H3K27ac (Abcam, #ab4729) with the following optimizations: 5−10 million cells were crosslinked with 1% formaldehyde at room temperature for 10 minutes. Prior to restriction digestion, sodium dodecyl sulfate (SDS) treatment was done at 62 °C for 10 minutes. Restriction digestion with MboI (New England Biolabs France, R0147M) was performed for 2 hours at 37 °C. Before the fill-in reaction, MboI was heat-inactivated at 62 °C for 10 minutes, followed by two washing steps of pelleted nuclei using 1x fill-in reaction buffer. After the fill-in reaction, ligation was carried out at 4 °C for 16 hours, ensuring effective proximity ligation for subsequent steps.

### HiChIP sequencing and data analysis

HiChIP libraries were sequenced in paired-end mode using the Illumina HiSeq 2500 system, achieving a depth of over 100 million read pairs (2 x 100 bp). HiChIP data analysis was done as describe previously [12]. HiChIP-seq paired-end reads were aligned to the human genome (hg38), duplicate reads were removed, reads were assigned to MboI restriction fragments, filtered into valid interaction pairs, and the interaction matrices were generated using the HiC-Pro pipeline default settings [91]. The config file of the HiC-Pro was set to allow validPairs at any distance from each other. HiC-Pro valid interaction reads were then used to detect significant interactions using: (1) Tag directories were created using the function makeTagDirectory from HOMER using the alignment BAM files from HiC-Pro output. The command line is (makeTagDirectory Tag R1.bam R2.bam -tbp 1). (2) We used runHiCpca.pl function from HOMER (http://homer.ucsd.edu/homer/interactions2/HiCpca.html) to perform the chromatin compartment analysis (PCA) of the data with the command (runHiCpca.pl -res 30000 -genome hg38 -cpu 8). (3) We used analyzeHiC from HOMER [92] to identify the HiChIP interactions (http://homer.ucsd.edu/homer/interactions/) as follows analyzeHiC Tag - res 10000 -cpu 8 -interactions -center -pvalue 0.05 -nomatrix. (4) And the interactions were annotated to the human genome (hg38) using the function annotateInteractions.pl from HOMER (http://homer.ucsd.edu/homer/interactions/HiCannotation.html). (5) HiChIP interaction hubs were generated using the analyzeHiC tool, and the interactions were annotated to the human genome (hg38) with the annotateInteractions.pl script, as described above. The interactions were extracted in BEDPE format to determine the total number of interactions, chromosomal distribution, and loop size. The loop size was represented as Log2 (size in kilobases). (6) The annotated HiChIP interactions were filtered by (a) the peaks of H3K27ac, then (b) with the reference of 746 upregulated genes, and (c) with the peaks of CTCF. The filtered loops were extracted in BEDPE format and then visualized using IGV genome browser. (7) The HiChIP contact matrices were generated using the allValidPairs output from HiC-Pro and its utilities (https://github.com/nservant/HiC-Pro/blob/master/doc/UTILS.md). Visualization of contact matrices was performed using Juicebox (https://www.aidenlab.org/juicebox/) from Juicer tools [93] with the following command line (hicpro2juicebox.sh -i .allValidPairs -g hg38.chrom.sizes -j juicer_tools.2.20.00.jar -r MboI_resfrag_hg38.bed) or with GENOVA package (https://github.com/robinweide/GENOVA) [94].

### Statistical Analysis

Depending on the dataset, various statistical tests were conducted to assess the significance of the results. The values of the statistical tests used across different experiments are available in the Source Data file. Additional details on the statistical analysis for specific experiments are provided in the figures and figure legends. Briefly, RNA-seq, ChIP-seq, CUT&Tag, and HiChIP samples were analyzed using next-generation sequencing. For other experiments presented here, samples were analyzed in triplicates, and experiments were performed three times. Statistical analyses were carried out using Excel Solver and using GraphPad Prism 8 software. Data displayed in box plots represent a five-number summary. One-tailed or two-tailed t-tests were employed to assess the differences between groups, with significance levels denoted as follows: \**P* ≤ 0.05; \*\**P* < 0.01; and \*\*\**P* < 0.001.

### List of tools used for analysis

Online tools and open-source software were used for analysis and to create figure panels as indicated. The Cancer Genome Atlas (TCGA) (https://www.cancer.gov/tcga) [95]; TCGAbiolinks [96]; Tumor suppressor gene database (<u>TSGeneHome</u>); KEGG database; ShinyGO 0.82 ([ShinyGO 0.82](https://bioinformatics.sdstate.edu/go/)); Juicebox Web App (Juicebox); MEME Suite [97]; JASPAR database ([JASPAR](https://jaspar.elixir.no/)); eRNAbase [73]; cBioPortal [98]; GWAS catalog [99]; KM plotter [74]; Microsoft Office Suite; R (https://www.r-project.org/); and GraphPad Prism 8.

## Data availability

The data that support this study are provided with this paper. Source data are provided with this paper as a Source Data file. The sequencing data generated in this study have been deposited in NCBI’s Gene Expression Omnibus database [100] under accession numbers GSE29ABC, GSE29DFG, GSE29HIJ and GSE29KLM. Furthermore, we retrieved and used publicly available datasets to aid analysis of our data. Supplementary Data 1 contains all data sets used in this study.

## Acknowledgments

We thank Roswitha Bender and Anne Robert for technical support; Diana G. Rogel-Ayala, Hector A. Aguirre-Alarcon, Karla Rubio, Virgine Marchand, Iouri Motorine and the EpiRNA-Seq Core Facility for support with NGS-based methods; Kerstin Richter, Alessandro Ianni, Sylvain Maenner and Bruno Charpentier for reagents; Sandrine Gulberti, Hervé Kempf, Jean-Baptiste Vincourt, Sylvie Fournel-Gigleux, Mohamed Ouzzine, Catherine Bui, Lydia Barré and Nick Ramalanjaona for comments. GB was funded by the “Centre National de la Recherche Scientifique” (CNRS, France), “Délégation Centre-Est” (CNRS-DR6) and the “Université de Lorraine” (UL, France) through the initiative “Lorraine Université d’Excellence” (LUE) and the dispositive “Future Leader”. GD and JC are supported by the CRC 1366 (Projects A03, A06), the CRC 873 (Project A16), the CRC1550 (Project A03) funded by the DFG, the DZHK (81Z0500202) funded by BMBF, the Medical Faculty Mannheim of University of Heidelberg (90703207) and the Baden-Württemberg foundation special program “Angioformatics single cell platform”. GS was awarded a doctoral fellowship and received support from the DrEAM mobility grant episode 10^th^, both by the UL (France) through the initiative LUE. TB is supported by the Deutsche Forschungsgemeinschaft, Excellence Cluster Cardio-Pulmonary Institute (CPI), Transregional Collaborative Research Center TRR81, TP A02, SFB1213 TP B02, TRR 267 TP A05 and the German Center for Cardiovascular Research.

## Authors Contributions Statement

GS, SG and GB designed and performed the experiments; JG, TB and GD were involved in study design; GB and GS designed the study; GS, JC, GB and SG analyzed the data; GB, GS, JG, and JC wrote the manuscript. All authors discussed the results and commented on the manuscript.

