## Supplementary information for "Erlotinib induces 3D genome rearrangements in lung cancer cells activating tumor suppressor genes through FOXA2-bound Epromoters"

[†]Lead contact


**This PDF file includes:**

Supplementary Figures 1 to 7

**Other Supplementary Materials for this manuscript include the following:**

Source Data file 01 - This is an Excel file that contains the data for all the plots presented in the article, including the values for statistical significance and the implemented statistical tests.

Supplementary Table 1 – This is an Excel file that contains a list with the accession numbers of all published NGS data sets used in this manuscript.

a RNA-seq data quality

| Sample | Total reads | Mapped reads | % Mapped (hg38) |
|---|---|---|---|
| A549 DMSO rep1 | 41,017,480 | 38,884,571 | 94.8% |
| A549 DMSO rep2 | 48,040,513 | 45,532,798 | 94.78% |
| A549 Erlotinib rep1 | 47,969,504 | 45,551,840 | 94.96% |
| A549 Erlotinib rep2 | 47,997,161 | 45,501,308 | 94.8% |

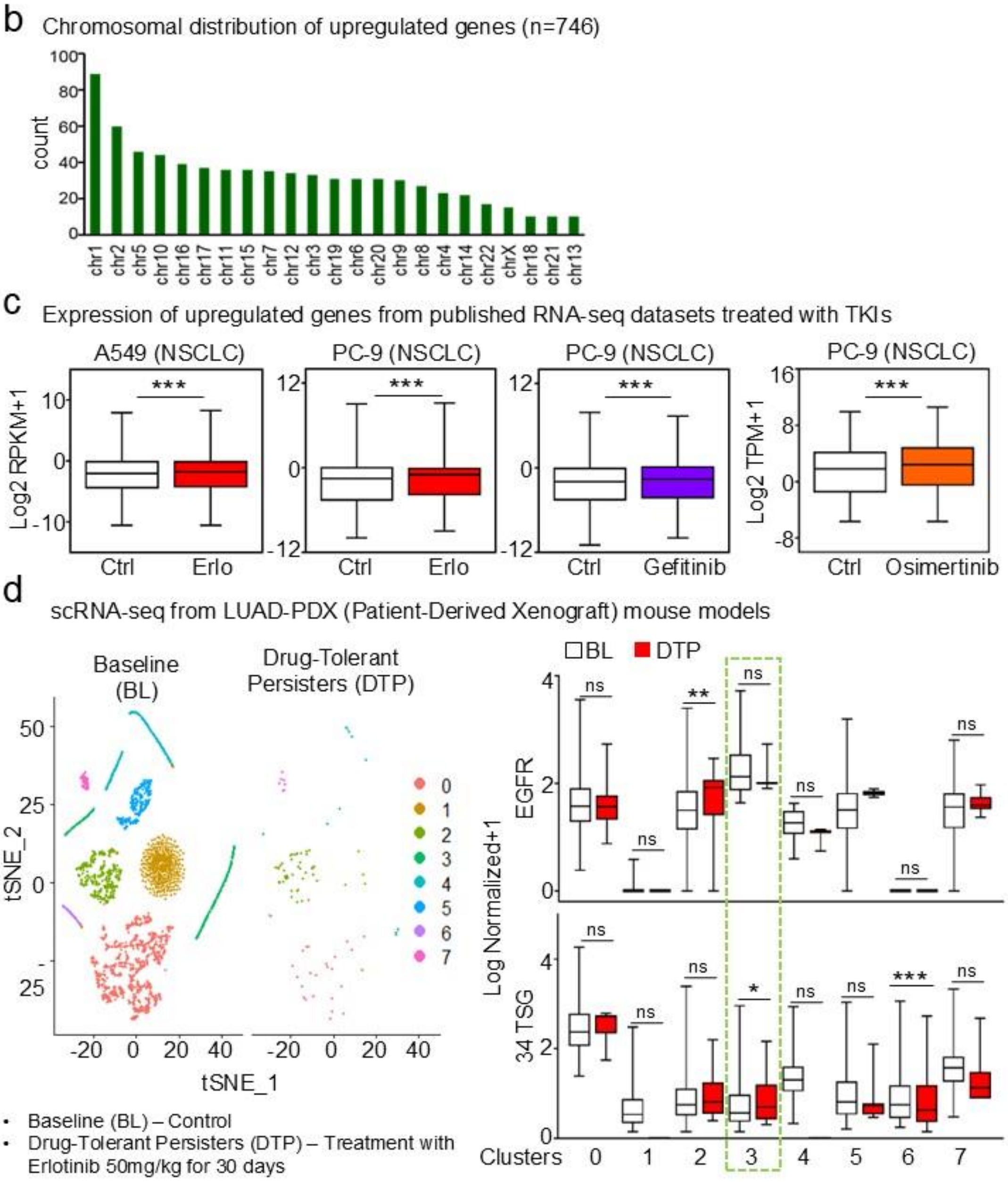


**Supplementary Figure 1: (a)** The description of the RNA-seq dataset supports the quality of the experiments. **(b)** Bar plot showing the distribution of upregulated genes (n=746) across all chromosomes. **(c)** Box plots of RNA-seq-based expression analysis of upregulated genes (n=746) from published datasets treated with different tyrosine kinase inhibitors (TKIs: Erlo; erlotinib, Gefitinib, and Osimertinib) in A549 and PC-9 cells. Values were normalized using RPKM, reads per kilobase of transcript per million mapped reads or TPM, transcript per million; represented as log2 RPKM + 1 or log2 TPM + 1. **(d)** Single-cell RNA sequencing (scRNA-seq) analysis of LUAD-PDX (Patient-Derived Xenograft) mouse models. The left panel displays a t-SNE plot comparing Baseline (BL) control cells and Drug-Tolerant Persisters (DTP) treated with erlotinib (50 mg/kg) for 30 days. Different clusters are labeled from 0 to 7. The right panel represents box plots with expression levels (Log Normalized+1) of EGFR and 34 tumor suppressor genes (TSG) across clusters for BL (white) and DTP (red) conditions. The green dashed box highlights cluster 3, showing significant differences in expression levels of EGFR and 34 TSG between BL and DTP conditions. All box plots display the median (middle line), 25th and 75th percentiles (box), and 5th and 95th percentiles (whiskers). Statistical significance is represented by asterisks: ***$P \leq 0.001$; **$P \leq 0.01$; *$P \leq 0.05$; ns, non-significant. *P*-values were calculated using two-tailed t-test or one-tailed Wilcoxon test (box plots). Source data are provided as a Source Data file.

a

H3K4me3 ChIP-seq data quality

| Sample | Total reads | Mapped reads | % Mapped (hg38) |
|---|---|---|---|
| Input DMSO | 13,536,624 | 11,235,397 | 83% |
| Input Erlotinib | 13,145,365 | 11,872,893 | 90.32% |
| H3K4me3 ChIP DMSO | 36,199,133 | 35,808,182 | 98.92% |
| H3K4me3 ChIP Erlotinib | 36,399,536 | 35,973,661 | 98.83% |

b

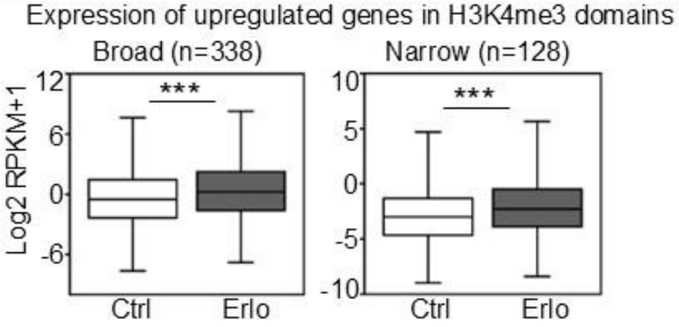


Supplementary Figure 2

**Supplementary Figure 2: (a)** The description of the H3K4me3 ChIP-seq dataset supports the quality of the experiment. **(b)** Box plots of RNA-seq-based expression analysis of upregulated genes (n=746) that were separated into the indicated H3K4me3 domains (broad; n=338 or narrow; n=128) in Ctrl or Erlo treated A549 cells. All box plots display the median (middle line), 25th and 75th percentiles (box), and 5th and 95th percentiles (whiskers). Statistical significance is represented by asterisks: ***$P \leq 0.001$; **$P \leq 0.01$; *$P \leq 0.05$; ns, non-significant. *P*-values were calculated using a two-tailed t-test (box plots). Source data are provided as a Source Data file.

a

H3K27me3 CUT&Tag-seq data quality

| Sample | Total reads | Mapped reads | % Mapped (hg38) |
|---|---|---|---|
| H3K27me3 CUT&Tag DMSO | 20,342,373 | 19,475,787 | 95.74% |
| H3K27me3 CUT&Tag Erlotinib | 20,799,413 | 20,063,113 | 96.46% |

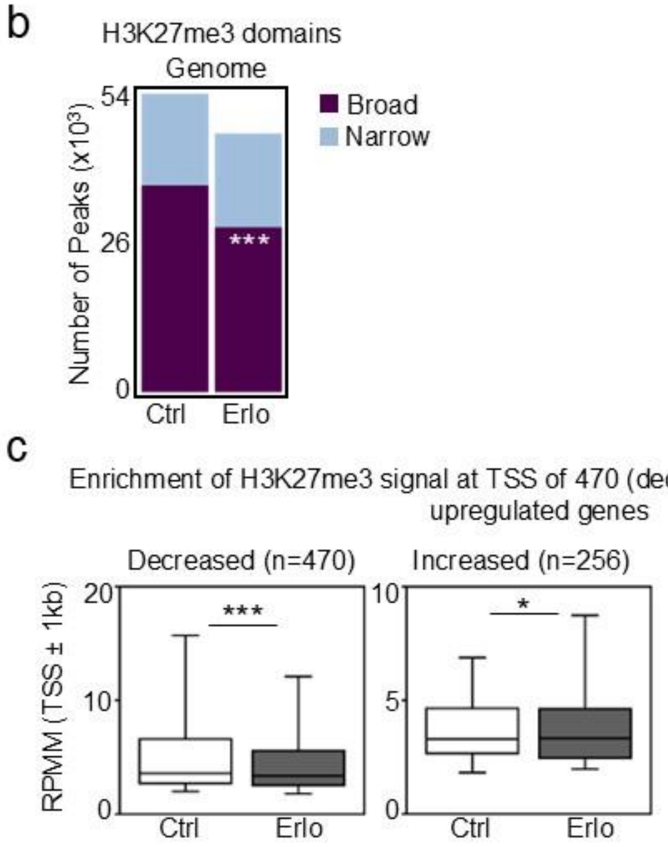


Supplementary Figure 3

**Supplementary Figure 3: (a)** The description of the H3K27me3 CUT&Tag dataset supports the quality of the experiments. **(b)** Bar plots displaying the broadness of H3K27me3 genome-wide in Ctrl or Erlo treated A549 cells. **(c)** Box plots showing the levels of H3K27me3 at the transcription start site (TSS -/+ 1 kb) of upregulated genes with decreased (n=470) and increased (n=256) H3K27me3 levels. Data were normalized using RPMM, read count per million mapped reads. All box plots represent the median (middle line), 25th, 75th percentile (box), and 5th and 95th percentile (whiskers). In all plots asterisks represent *P*-values, ***$P \leq 0.001$; **$P \leq 0.01$; *$P \leq 0.05$; ns, non-significant. *P*-values were calculated after two-tailed t-test (box plots) or two-tailed Fisher exact test (bar plots). Source data are provided as a Source Data file.

**a** H3K27ac ChIP-seq data quality

| Sample | Total reads | Mapped reads | % Mapped (hg38) |
|---|---|---|---|
| H3K27ac ChIP DMSO | 32,929,636 | 30,229,405 | 91.8% |
| H3K27ac ChIP Erlotinib | 41,213,459 | 37,916,382 | 92% |

**b** H3K27ac HiChIP-seq data quality

| Sample | Total reads (Paired-end) | Alignment_R1 (hg38) | Alignment_R2 (hg38) | Valid pairs | Invalid pairs |
|---|---|---|---|---|---|
| H3K27ac HiChIP DMSO | 115,670,036 | 95.00% | 93.30% | 79% | 21% |
| H3K27ac HiChIP Erlotinib | 134,459,958 | 94.90% | 92.60% | 84.10% | 15.90% |

**c** Expression of upregulated genes looping to SE or TYE

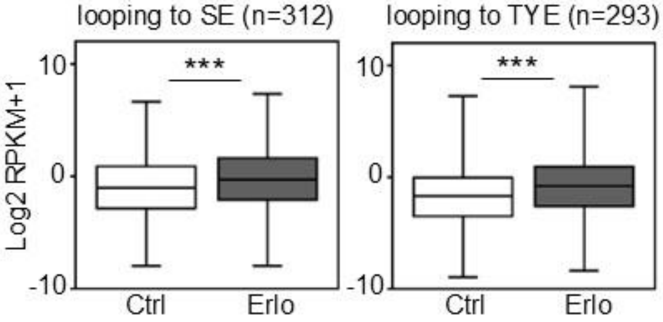


Supplementary Figure 4

**Supplementary Figure 4: (a)** The description of the H3K27ac ChIP-seq dataset supports the quality of the experiments. **(b)** The description of the H3K27ac HiChIP-seq dataset supports the quality of the experiments. **(c)** Box plots of RNA-seq-based expression analysis of upregulated genes that loop to super enhancers (SE; n=312) or typical enhancers (TYE; n=293) in Ctrl or Erlo treated A549 cells. All box plots display the median (middle line), 25th and 75th percentiles (box), and 5th and 95th percentiles (whiskers). Statistical significance is represented by asterisks: ***$P \leq 0.001$; **$P \leq 0.01$; *$P \leq 0.05$; ns, non-significant. *P*-values were calculated using a two-tailed t-test (box plots). Source data are provided as a Source Data file.

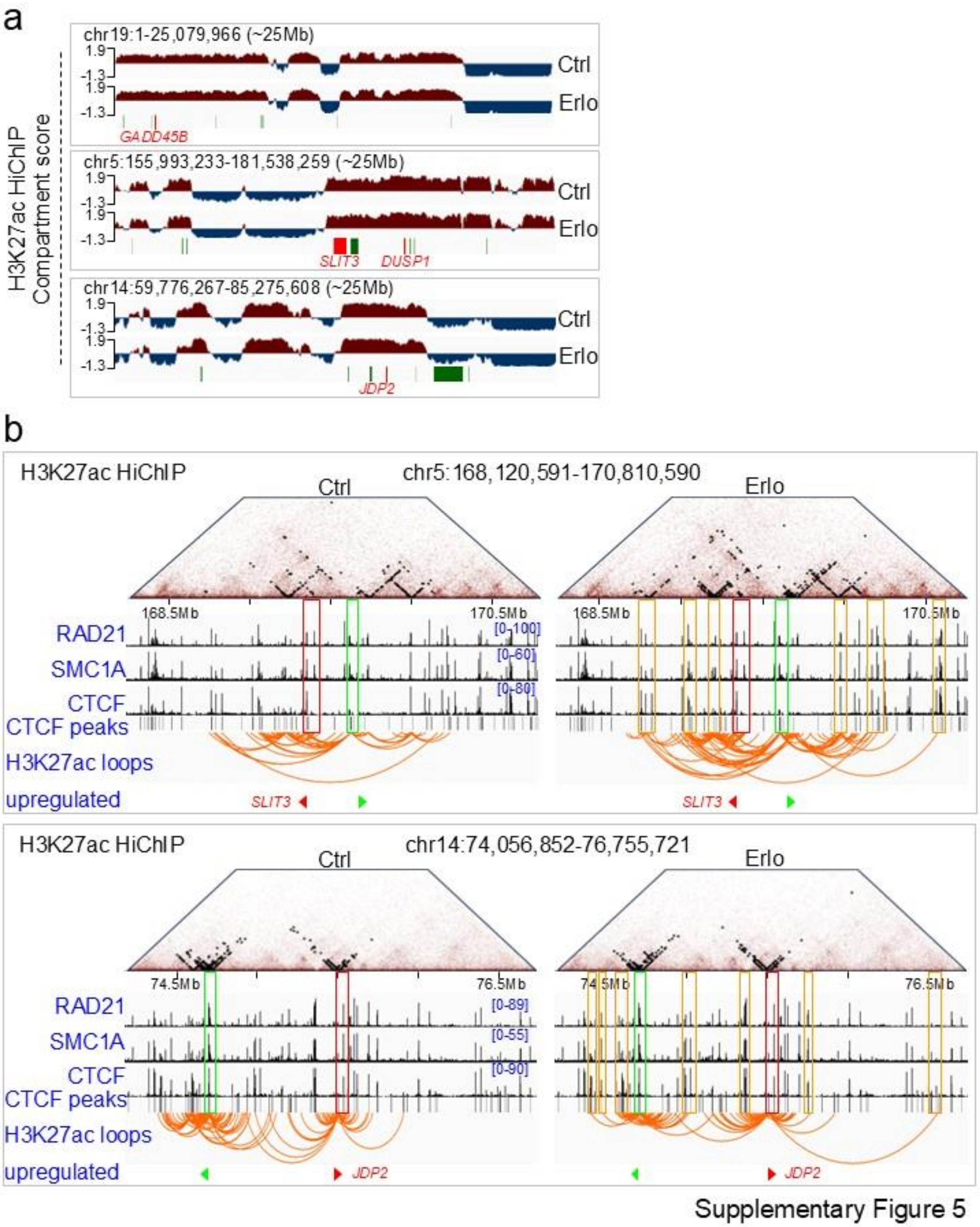


**Supplementary Figure 5: (a)** Changes in chromatin compartmentalization are visualized across a ~25Mb (Mb; megabases) window using compartment scores (y-axis) derived from H3K27acspecific HiChIP experiments in the Ctrl and Erlo treated conditions. Upregulated genes in clusters are highlighted in green, and key tumor suppressor genes (TSG) are highlighted in red (*GADD45B*, *SLIT3*, *DUSP1*, and *JDP2*). **(b)** Snapshots depict H3K27ac-specific chromatin contact matrices represented as pyramid plots comparing control (Ctrl, left) and erlotinib (Erlo, right) treated conditions at the top. Black dots within the pyramids represent the number of H3K27ac-specific interactions. Below the pyramid plots, ChIP-seq tracks illustrate factors of the cohesin complex (RAD21, SMC1A) and CTCF, along with CTCF peak annotations, and H3K27ac-specific chromatin loops. The visualization spans a ~2.5Mb window surrounding the promoters of key TSG (red triangle) and upregulated gene in clusters (bright green triangles). Co-enrichment of RAD21, SMC1A, and CTCF at TSG promoter is highlighted in red boxes, while upregulated genes are marked with bright green boxes. Regions looping to the promoters are indicated by orange boxes, illustrating chromatin interactions. The IGV genome browser was used for visualization.

a

ATAC-seq data quality

| Sample | Total reads | Mapped reads | % Mapped (hg38) |
|---|---|---|---|
| PC9 ATAC-seq DMSO | 30510312 | 24319770 | 79.71% |
| PC9 ATAC-seq Erlotinib | 31970207 | 27366497 | 85.60% |

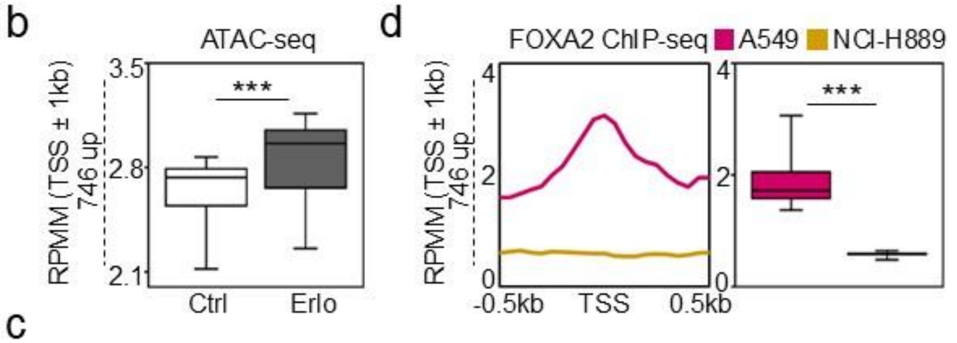


c

FOXA2 ChIP-seq data quality

| Sample | Total reads | Mapped reads | % Mapped (hg38) |
|---|---|---|---|
| NCI-H889 FOXA2 ChIP | 15509089 | 13674364 | 88.17% |
| NCI-H889 Input | 6528652 | 6414401 | 98.25% |
| FOXA2 ChIP from A549 cells were accessed from ENCODE (ENCSR000BRE) | | | |

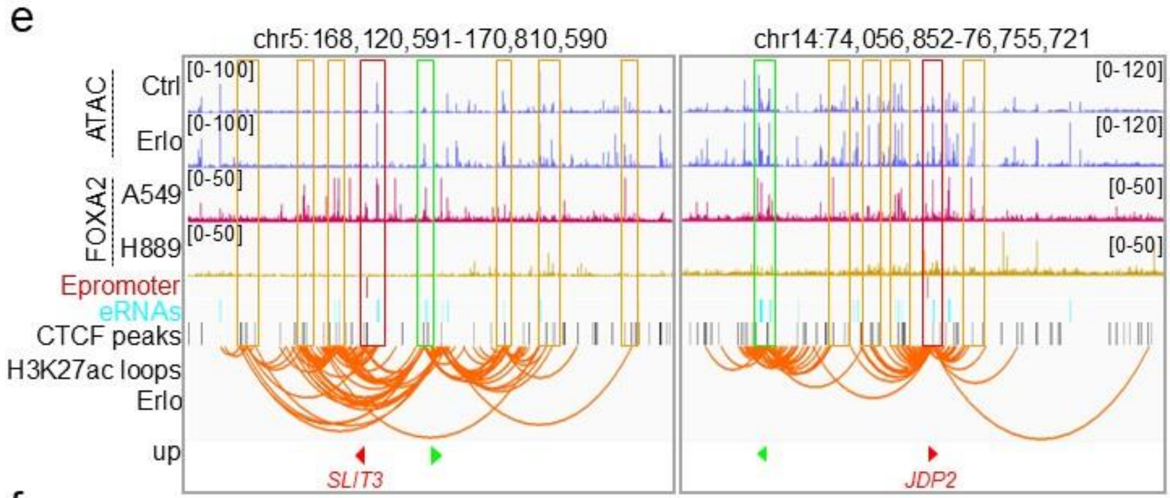


f

RNA-seq data quality

| Sample | Total reads | Mapped reads | % Mapped (hg38) |
|---|---|---|---|
| Ctrl rep1 | 8103189 | 7,315,559 | 90.28% |
| Ctrl rep2 | 7747186 | 7,053,038 | 91.04% |
| siFOXA2 rep1 | 14129418 | 12,732,018 | 90.11% |
| siFOXA2 rep2 | 22511259 | 19,843,674 | 88.15% |

Supplementary Figure 6

**Supplementary Figure 6: (a)** The description of the ATAC-seq dataset supports the quality of the experiments. **(b)** Box plots from ATAC-seq showing the accessibility at the transcription start site (TSS -/+ 1 kb) of 746 upregulated genes in Ctrl or Erlo treated PC-9 cells. Data were normalized using RPMM, read count per million mapped reads. **(c)** The description of the FOXA2 ChIP-seq dataset supports the quality of the experiments. **(d)** Aggregate and box plots showing the FOXA2 ChIP-seq enrichment at transcription start site (TSS -/+ 1 kb) of 746 upregulated genes in A549 and NCI-H889 cells. **(e)** Genome browser snapshots of selected tumor suppressor genes (TSG) loci showing chromatin accessibility by ATAC-seq (blue) in Ctrl and Erlo treated PC-9 cells, FOXA2 ChIP-seq in A549 (magenta) and NCI-H889 cells (gold). The tracks below depict Epromoters (red), eRNAs (arctic blue), CTCF peaks (black), and H3K27ac-specific chromatin loops in Erlo treated A549 cells. The visualization spans a ~2.5Mb window surrounding the promoters of key TSG (red triangle) and upregulated gene in clusters (bright green triangles). Co-enrichment of FOXA2 with ATAC-seq, Epromoter, eRNAs, and CTCF peaks at TSG promoter is highlighted in red boxes, while upregulated genes are marked with bright green boxes. Regions looping to the promoters are indicated by orange boxes, illustrating chromatin interactions. The IGV genome browser was used for visualization. **(f)** The description of the RNA-seq dataset supports the quality of the experiments. Source data are provided as a Source Data file.

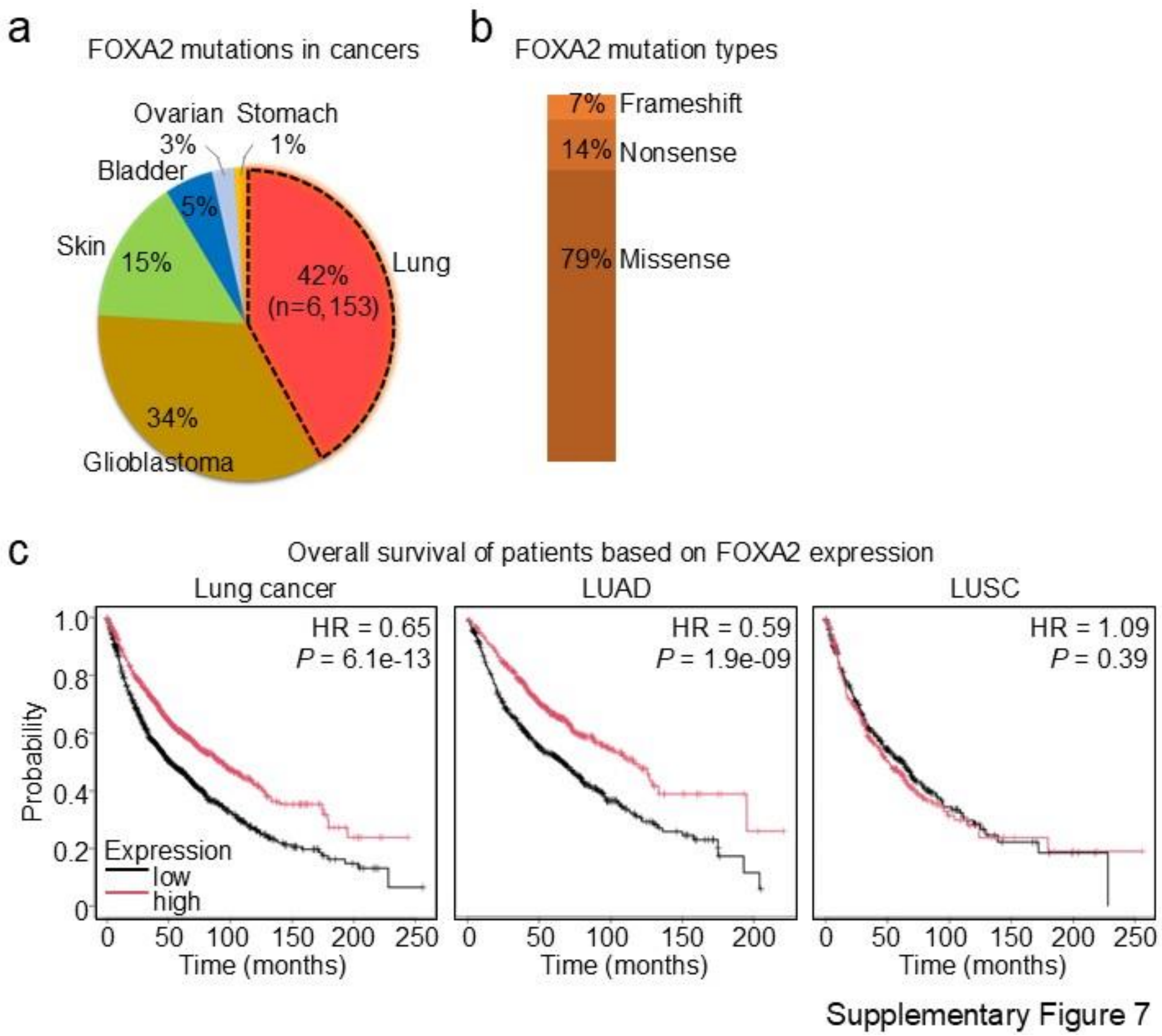


**Supplementary Figure 7: (a, b)** Pie chart showing the single nucleotide polymorphisms (SNPs) reported as FOXA2 mutations in various cancers. The segments represent different cancers, with lung cancer showing the highest proportion of FOXA2 mutations (42%, n=6,153), followed by other cancer types. In **(b)**, bar plot illustrates the types of FOXA2 mutations in lung cancer, the mutation categories include frameshift, nonsense, and missense as indicated. **(c)** Kaplan-Meier survival curves illustrating overall survival of patients based on the expression of FOXA2 across different lung cancer types. Survival probabilities are represented for all lung cancer patients (left), LUAD patients (middle), and LUSC patients (right). Patients with high TSG expression (red curves) show distinct survival patterns compared to those with low expression (black curves).

Lung cancer (HR = 0.65, *P* = 6.1e-13, median survival in months: high expression vs. low expression is 92.97 vs. 51), LUAD (HR = 0.59, *P* = 1.9e-09, median survival in months: high expression vs. low expression is 116 vs. 66), and LUSC (HR = 1.09, *P* = 0.39, median survival in months: high expression vs. low expression is 51 vs. 62.47). HR, hazard ratio. Source data are provided as a Source Data file.

**Data S1. (separate files)**

Source Data file 01 - This is an Excel file that contains the data for all the plots presented in the article, including the values for statistical significance and the implemented statistical tests.

Supplementary Table 1 – This is an Excel file that contains a list with the accession numbers of all published NGS data sets used in this manuscript.
